# Expanding the definition of MHC Class I peptide binding promiscuity to support vaccine discovery across cancers with CARMEN

**DOI:** 10.1101/2025.05.08.651510

**Authors:** Michal Mateusz Waleron, Ashwin Adrian Kallor, Aleksander Palkowski, Emilia Daghir-Wojtkowiak, Piyush Borole, Mikolaj Kocikowski, Karol Polom, Fabio Marino, Beatriz Monterde, Michele Mastromattei, Davide Venditti, Mark Stares, The KATY Consortium, Ashita Singh, Theodore Hupp, Christophe Battail, Alexander Laird, Catia Pesquita, Luis Zapata, Stefan N. Symeonides, Ajitha Rajan, Fabio Massimo Zanzotto, Javier Antonio Alfaro

## Abstract

Promiscuity in T-cell antigen landscapes refers to the dual flexibility of peptides binding multiple MHC alleles and MHC alleles presenting diverse arrays of peptides. By understanding how neoantigens are shared across varied HLA backgrounds, promiscuity analysis can inform the selection of cancer-vaccine targets that reach a wider segment of the population and help refine patient stratification for diverse immunotherapies. We expand the concept of promiscuity to encompass peptides, MHC alleles, individuals, populations, and genomic regions. Our CARMEN database release harmonizes data from 72 publications (2,323 samples) across tissue types, with a focus on cancer. Using Gibbs clustering and dimensionality reduction (UMAP), we systematically map promiscuity and immunological versatility across these biological levels. Gene and mutation analysis reveals recurrent cancer mutations in highly promiscuous genomic regions, highly mutated cancer genes that avoid presented regions, and sheds light on genomic regions important to response to immunotherapy.

## Introduction

Cancer vaccines and immunotherapies, including peptide-based, viral, dendritic cell, and mRNA vaccines, as well as checkpoint inhibitors, cellular therapies, bispecific T-cell engagers, and other emerging modalities are advancing cancer therapy^1–5^. Recent trials with mRNA vaccines for pancreatic cancer^6^, melanoma^7^, and glioblastoma^8^ highlight their promising clinical activity. Understanding the MHC-I T-cell response and the patient populations that would display a satisfactory response is critical for neoantigen selection and immunotherapy response prediction. Immunopeptidomic Mass Spectrometry has enhanced antigen presentation analysis, improving personalized immunotherapy strategies^9,10^.

Two foundational concepts in antigen presentation studies are promiscuity — where peptides can bind multiple MHC alleles — and hotspots, referring to genomic regions consistently presented to the immune system. Studies have shown that peptide binding breadth influences immune recognition^11,12^, and other research has demonstrated broader binding in contexts of low MHC expression^13^. Other studies have identified promiscuous HER2/neu epitopes^14^ and explored the ability of MHC class I molecules to bind millions of extracellular peptides with high affinity while maintaining selective presentation to T cells^15^. Studies have started to report that hotspots enhance immune surveillance and may serve as key immunotherapy targets, especially when overlapping mutations are present^16,17,18^.

With MHC Class I binding promiscuity as a guiding principle, we developed the CAnceR iMmunopEptidogeNomics (CARMEN) database. By mapping physically detected MHC Class I peptides to a novel motif atlas, we make clear the small fraction of anchor-residue combinations accounts for most peptides, indicating a surprisingly simple but continuous antigen-binding landscape. We expand “promiscuity” beyond the classic peptide–MHC perspective to also encompass individuals, populations, and genomic regions, uncovering numerous “hotspots” with high immune visibility and recurrent cancer mutations. Some well-known oncogenes (e.g., KRAS, TP53) contain highly promiscuous continuous genomic regions (contigs) covered by the immunopeptidome, whereas other frequently mutated genes remain less immunologically visible.

We focus on understanding promiscuity from the physical evidence of what is actually detected by mass-spectrometry as opposed to prediction. To bridge genomics and immunopeptidomics, we redefine antigen presentation hotspots through epitope contigs and scaffolds, tracing contiguous genomic regions presented in the immunopeptidome. We systematically map peptides to the genome, keeping track of promiscuity, unique peptide counts, expression, and MHC-I allele associations. Reanalyzing 72 public immunopeptidomic datasets, we identified 816,222 unique peptides, consolidated into CARMEN: Cancer ImmunoPeptidogenomics, linking canonical and cryptic peptides to their genomic origins. Motif analysis of all identified peptides revealed a reduced complexity of antigen binding space and allele frequency overlaps across populations.

Integrating these promiscuity metrics at the peptide and contig levels enhances the prediction of immunotherapy response in clear cell renal cell carcinoma. To assess clinical relevance of genomic regions observed in the immunopeptidome, we analyzed 311 renal cancer patients from Braun *et al*.^19^ using an SVM model on contig-overlapping mutations. Some contigs correlated with highly immune-visible regions, suggesting their value for predicting immunotherapy response and in guiding vaccine design. In short, we formalize a broader concept of promiscuity, positioning CARMEN as a comprehensive resource for peptide, MHC allele, contig, scaffold, and population-level analysis of diversity in antigen presentation.

## Results

The CARMEN database provides a comprehensive, pan-cancer view of antigen presentation, identifying highly immune-visible genomic regions attractive for immunotherapy (**Figure 1a-e**). It spans 31 cancer types from 2,323 samples, comprising 1,400 cancerous, 849 healthy, and 41 benign samples (**Figure 1e)**. Using previously published approaches^20^, we re-analyzed 72 immunopeptidomic mass spectrometry datasets (2015–2023), covering 184 experimentally determined HLA alleles (**Figure 1d**, **Supplementary Table 1**). Sample sources include patient-derived tissues (1,169, 50%), cell lines (910, 39%), and primary cells (244, 11%) (**Figure 1e**). 1,607 samples had published experimentally determined MHC Haplotype.

**Figure 1:**
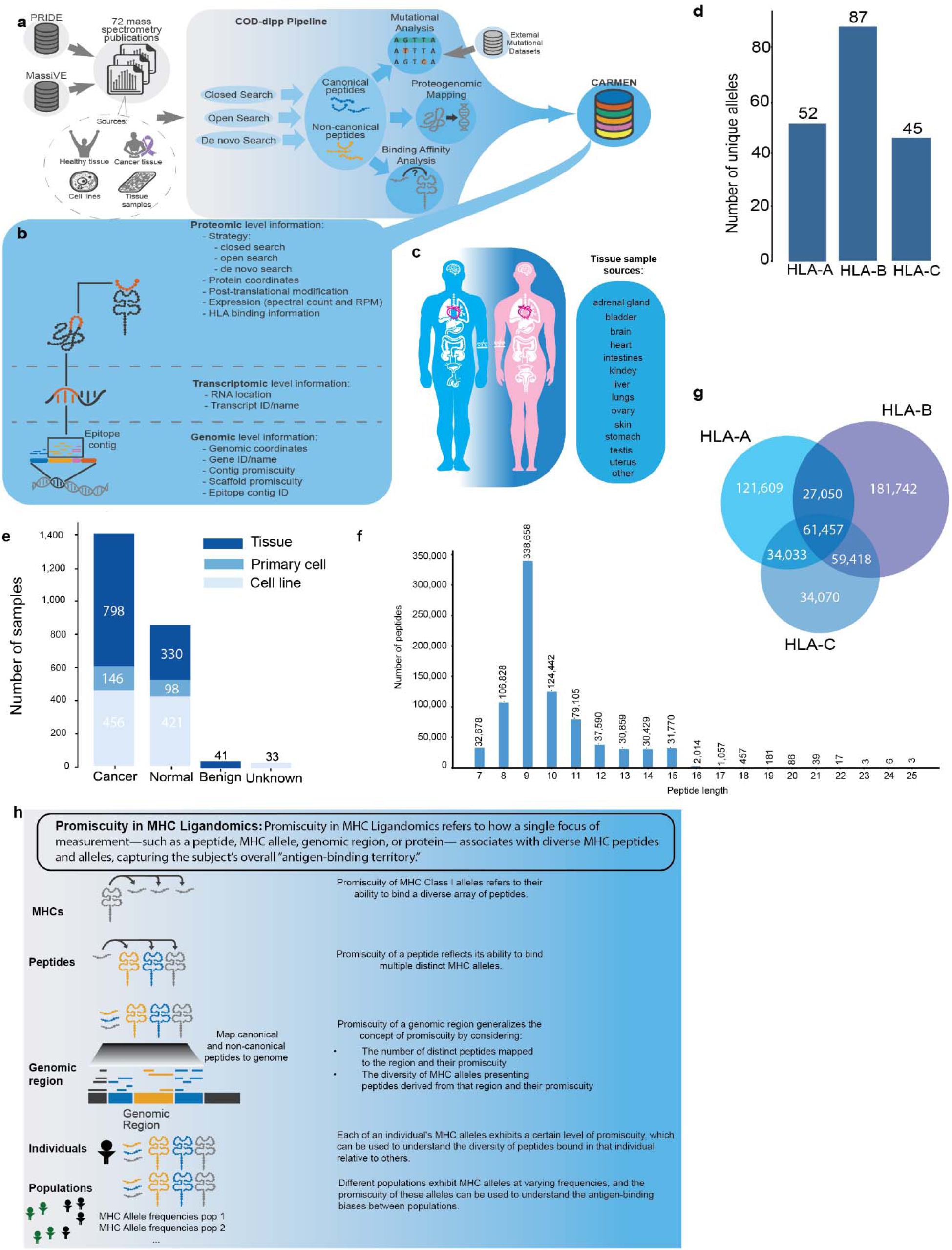
CARMEN, a proteogenomic database defining physically detected promiscuity and antigen recurrence in cancer. At its core, CARMEN integrates MHC Class I peptide data from 72 immunopeptidomic mass spectrometry publications (MASSIVE and PRIDE), featuring **(a)** the COD-Dipp pipeline’s search workflows—closed, open, and de novo—and **(b)** a proteogenomic mapping framework, forming a multi-omic resource with layers spanning the central dogma. **(c)** The dataset is obtained from 21 different tissues which could be cancerous, healthy or benign**. (d)** All three major classes of HLA alleles are represented in the CARMEN database. **(e)** Overall, the CARMEN database comprises normal, cancerous and benign samples taken from tissues, cell lines and primary cells, with cancer tissues comprising the largest share of samples. **(f)** The peptide lengths show the expected peak at 9 amino acids and the bulk of the distribution observed between 7 to 11 amino acids. **(g)** NetMHCpan-4.1 was used to predict peptide binding to the experimentally determined MHC alleles from the samples, illustrating the predicted overlap of peptides across the three alleles. **(h)** This resource formalizes promiscuity in MHC ligandomics, describing it for any set of peptides across hierarchical levels: individual peptides, MHC alleles, genomic regions, individuals, and populations.

The database contains 816,222 unique peptides, categorized as canonical (797,561), non-canonical (8,743), and mapping to both (9,918). The length of most peptides (95%) is within the range of 8–12 amino acids, with 70% (569,928) being 8–10 amino acids long and a mode of 9 amino acids (338,658, 41%) (**Figure 1f**). A minority of peptides over 15 amino acids may represent HLA-II binders or other contaminants, despite all samples originating from MHC Class I studies. Samples were of varying quality. The median peptide count per sample was 2,052, at the other extreme 44 samples yielded only one peptide under our rigorous pipeline (**Supplementary Table 2**). For each peptide, its sample-related MHC(s) of origin was checked by NetMHCpan-4.1. This prediction suggests a concentration of strong binders for HLA-B (n = 329,667), followed by HLA-A (n = 244,149) and HLA-C (n = 188,978). 61,457 peptides appeared to be shared between all the three classes, while a minority of peptides was unique to a single HLA class (**Figure 1g**). However, the robustness of prediction-based data may be uncertain, especially in case of HLA-C where data may be limiting. CARMEN helps to clarify what is observed by physical detection and cross-presentation as opposed to prediction.

### Data Processing & Mass Spectrometry Searches

We downloaded and reprocessed 72 datasets from PRIDE and MASSIVE, selected using Bedran *et al.’s* ^20^ criteria for sample type, mass spectrometry technology (LTQ Orbitrap), experiment type (immunopeptidomics), and acquisition method (DDA) (**Figure 1a–c**). Raw files were converted to mzML using MSConvert, and a minimum of 25-peptide identifications was set to control for the sample quality variability observed in **Supplementary Table 2**.

We used COD-Dipp^20^ identically to the original publication for mass spectrometry searches, integrating MSGF+ (closed searches), MS-Fragger (open searches), and DeepNovo V2 (de novo searches) (The search parameters used to perform the closed and open searches are detailed in **Supplementary File 1** and **Supplementary File 2** respectively). Identified peptides were mapped to hg38.p13 using Pogo^21^, with genomic coordinates exported as BED files for interoperability with genomic pipelines. NetMHCpan-4.1^22^ annotated peptides based on MHC-I binding affinity, classifying them as strong (SB ≤ 0.2), weak, or non-binders (WB ≥ 1.2) based on experimental sample haplotypes where known (**Supplementary Table 3**). The individual search results were then engineered into the CARMEN dataset.

### Engineering the CARMEN dataset

We engineer and release the CARMEN dataset^23^, a proteogenomic repository of immune peptides presented in both normal and cancer tissues and disseminated it adhering to F.A.I.R. principles as part of the European Open data pilot. It consists of the following four tables:

1. *carmen-main.parquet*: The main database table, consisting of primary data related to each peptide identification, including their sample and experiment of origin, sequence, type (canonical/non-canonical/indeterminate) promiscuity and binding HLA allele.
2. *carmen-mapped-protein-annotations-msfragger.parquet*: The proteomic data associated with each peptide, including the peptide sequence, its gene/transcript/protein of origin and its protein start and end points.
3. *carmen-mapped-protein-annotations-pogo.parquet*: The genomic data associated with each peptide, including the peptide sequence and genomic coordinates obtained from the PoGo algorithm.
4. *carmen-netmhcpan-scores.parquet*: A table of the binding affinity scores of all the peptides in our database scored against every known HLA allele available in the NetMHC allele database.

A more detailed description of each table along with their associated metadata is found in the file titled **Readme.md**.

### Deriving and visualising the MHC Class I motif atlas in CARMEN

MHC Class I peptides are typically thought to have two conserved sites known as anchor residues. We analyzed 9-mer sequences in CARMEN to see how residues are conserved, to map out the space of total possible combinations of residues with x-sites of conservation given by

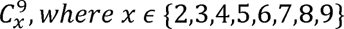

Exploring this entire space for different values of x, we characterized the optimal number of amino-acid combinations at specific sites along the peptide that could explain the largest fraction of peptides in the CARMEN dataset (**Figure 2a**). For 2 positions of conservation we found that 90% of peptides could be grouped into just 483 amino acid patterns. We release the results for the other combinations as a part of the CARMEN database (**Supplementary Figure 1**). Overall, the new CARMEN release extends detections beyond the MHC-Motif Atlas^24^. As shown in **Figure 2b** CARMEN extends the 9-mer peptide landscape by 196,869 peptides offering a deeper interrogation of MHC Class I binding patterns.

**Figure 2:**
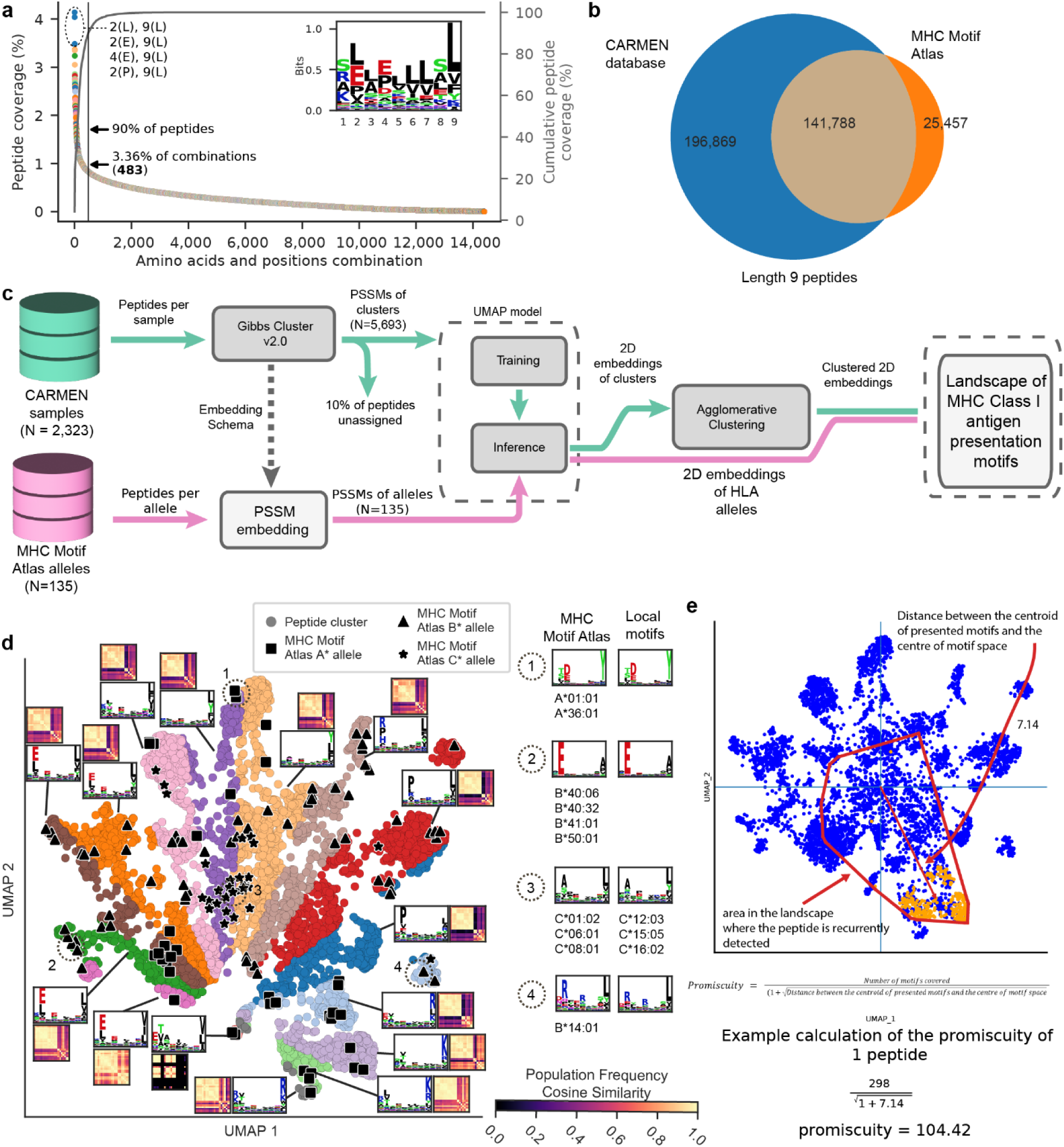
Defining promiscuity of physically detected peptides in immunopeptidomics datasets. Focused on 9-mer peptides, we explored peptide recurrence in the physically identified immunopeptidome by analysing **(a)** The fraction of 9-mers in the dataset explained by specific patterns in the peptides. I.e amino acid A at position k and amino acid B at position j. **(b)** The overlap of peptides identified in CARMEN and those identified in the HLA motif atlas. **(c)** Clustering peptides across all samples using Gibbs cluster plus and visualizing the similarities in the motifs defining these clusters alongside those of the HLA motif atlas by UMAP to produce a **(d)** landscape of antigen binding patterns in the immunopeptidome. The numbered circles demonstrate that different local neighbourhoods of MHC motif atlas^24^ alleles are grouping similar motifs. The colours on the UMAP partition the space into 13 “***broad clusters***”, with motifs shown for the entire region surrounding the figure. **(e)** Finally, using the revealed UMAP map we developed a measure of ***promiscuity*** that includes the number of gibbs clusters recurrently presenting a peptide from a peptide list, and the broadness of the territory in the UMAP covered by clusters (defined by the magnitude of the vector to the centroid of those clusters). Exemplified in (d) is the promiscuity calculation for a single peptide that occupies a specific territory in the revealed landscape. Motifs are generated using the logomaker package (39).

Immunopeptidomics datasets usually blend peptides from several co-expressed HLA-I alleles, masking allele-specific motifs and hindering global analyses of HLA diversity and promiscuity. To disentangle this complexity across 1000s of samples, we applied an unsupervised clustering-and-embedding strategy that converts each mixed sample into a set of comparable motif signatures. An HLA motif landscape of CARMEN peptides was revealed using GibbsCluster^25^ (**Figure 2c**) and visualized by UMAP. We applied Gibbs clustering to generate Position Specific Scoring Matrices (PSSMs) for peptides across 2,323 CARMEN samples, followed by UMAP projection to visualise HLA binding similarities. Peptide sequences were clustered sample-wise, starting with the peptide list for each sample and using the Blosum62 substitution matrix, assuming 3-7 clusters based on 3-6 MHC alleles per sample. The result was a list of Position Specific Scoring Matrices (PSSMs) for each sample. The optimal number of PSSMs was determined using the Kullback-Leibler distance (KLD), with the highest KLD indicating the optimal number of clusters for each sample, yielding varying numbers of PSSMs per sample. UMAP was used to visualize the PSSM landscape and reduce data dimensionality (50 neighbours, minimum distance 0.5, cosine metric) (**Figure 2d**). 2D embeddings from UMAP were then input into an Agglomerative clustering algorithm, with a predefined number of clusters – termed “***broad clusters***”. For each “***broad cluster***” we summarize the motif change across the map by generating a motif from the peptides present in that part of the landscape. We also show that local-neighbourhoods in this UMAP visualization conserve HLA-motif atlas^24^ motifs by showing that motif logos for peptide clusters in the local neighbourhood of an HLA-motif atlas^24^ cluster resemble each other which fall into clear territories in the map.

Overall, CARMEN defines a limited yet continuous motif-atlas across our sample set (**Figure 2d**), consistent with the earlier combinatorial analysis (**Figure 2a**). While four distinct motifs appear to represent the extremes of this space, the samples contain motifs that blend seamlessly across the continuum.

### Defining promiscuity using the CARMEN MHC Class I binding motif atlas

We define promiscuity in terms of the ability of a collection of peptides to be found throughout the motif landscape revealed in **Figure 2d**, visualized for 1 peptide in **Figure 2e**. This definition applies universally to any set of peptides, for example representative of an MHC allele, an individual, a population, or genomic region of interest after mapping peptides to the genome (**Figure 1h**). Promiscuity reflects how widely a set of peptides is recurrently found across the UMAP landscape of Gibbs clusters (motifs) of physically detected peptides, indicating the experimental diversity of MHC Class I binding interactions.

To quantify promiscuity, we develop a promiscuity score based on any given list of peptides by analysing their distribution and recurrence within the UMAP landscape. Specifically, the promiscuity score captures both the breadth of motifs covered by the peptides and the diversity of these motifs within the overall motif space:

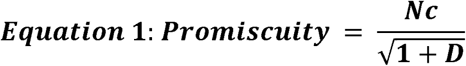

- **Number of Motifs Covered (*Nc*)**: This represents the count of distinct motifs that the peptides are associated with in the UMAP landscape.
- **Distance between the Centroid of Presented Motifs and the Center of Motif Space (*D*)**: This measures how far the collective set of presented motifs is from the central point of the entire motif space, reflecting the diversity and uniqueness of the motifs. We found that local-neighbourhoods commonly presented the same peptide and this term dampens the relative importance of the local neighbourhood, which would increase the magnitude of this vector.

By combining these two components, the promiscuity score captures not just the quantity of different motifs the peptides bind to but also the diversity of those motifs within the broader human motif landscape. A higher promiscuity score indicates that the peptides are not only binding to a larger number of motifs but are also interacting with motifs that are more diverse and spread out within the motif space. This metric can be applied across various biological levels by grouping peptides together and assessing their promiscuity. We demonstrate the promiscuity score calculation of a single peptide in **Figure 2e**.

### Exploring promiscuity in different contexts

#### Peptide promiscuity: recurrence of peptides across the CARMEN MHC Class I motif atlas

While studies such as in the development of SOPRANO by Zapata et al.^26^ have used netMHCpan-based predictions to map antigen presentation promiscuity across common and patient-specific HLA alleles, these models rely on computational inference of binding rather than direct measurement. In contrast, our approach leverages physical detection of MHC-bound peptides to empirically examine the cancer immunopeptidome. This comparison offers a valuable opportunity to assess how well predictive models reflect the complexity and specificity of antigen presentation in actual tumour contexts.

We provide a table of peptide-level promiscuity in **Supplementary Table 6** and a histogram of calculated promiscuities for each peptide in CARMEN in **Figure 3a**, clearly illustrating the fraction of overlap between experimental and SOPRANO-predicted peptides identified considering the most frequent 75 MHC Class I alleles in the human population. While the SOPRANO criteria—peptides predicted to bind any of the most frequent 75 MHC Class I alleles—do not directly quantify promiscuity, we identified a CARMEN promiscuity threshold of 13, below which concordance between predicted and experimentally detected peptides declines. Figures 3b and 3c show the overlap between physically detected peptides from CARMEN and SOPRANO predictions, both before and after applying this promiscuity cutoff. Applying the threshold reveals that CARMEN expands the known promiscuous immunopeptidome landscape by 55%, in terms of unique peptides, compared to the prediction methods used in Zapata et al. ^26^ (**Figure 3 b,c**).

**Figure 3:**
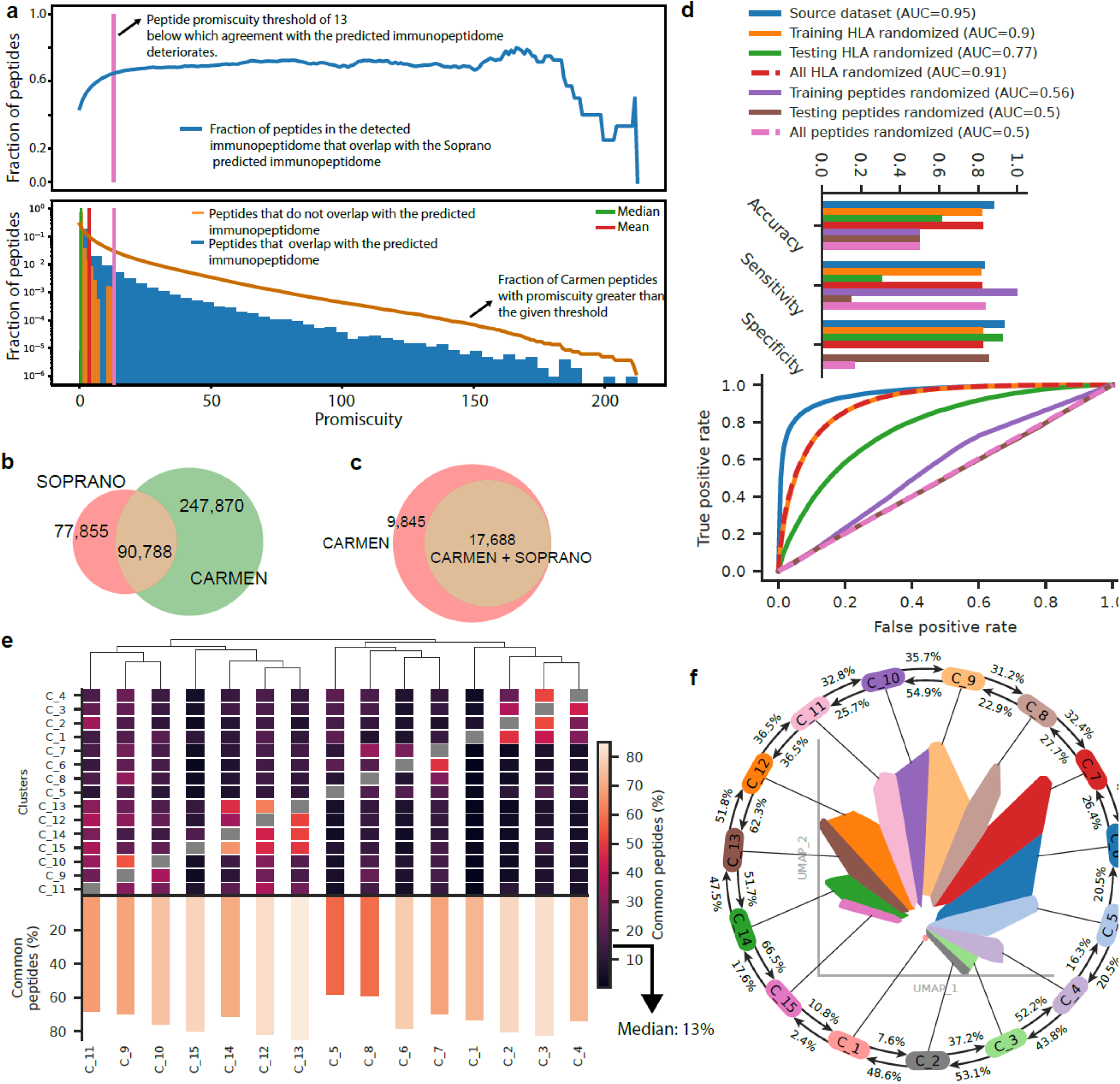
Promiscuity of peptides reveals the simplicity of the immunopeptidome. **(a)** Histogram of peptide promiscuities from the CARMEN database, illustrating the overlap between experimental and SOPRANO-predicted peptides across the 75 most common human MHC Class I alleles. A cut-off point is shown to illustrate our proposed definition of high-promiscuity based on agreement between experimental and predicted immunopeptidomes. **(b)** Venn diagram of predicted and experimentally detected unique peptides mapped to genomic regions in CARMEN vs SOPRANO. **(c)** Effect of applying a promiscuity threshold of 13 to the overlap between SOPRANO and CARMEN peptide-mapped genomic regions. **(d)** Predictive models can tell whether a peptide can be presented somewhere in the immunopeptidome with high accuracy, completely disregarding sample-specific MHC Allele information. **(e,f)** Exploring peptide recurrence across different parts of the revealed UMAP landscape. **(e)** Heatmap and histogram depict overall recurrence of peptides between clusters in the UMAP. Overall, one region of the UMAP has on average 15% overlap to any other, and adjacent regions **(f)** have very significant recurrence in peptides identified.

Another way to look at promiscuity in the revealed landscape is to understand the extent to which the overlap of peptides across different regions in the revealed motif map. We used a hierarchical clustering algorithm with the cosine similarity metric to split the UMAP atlas into regions we call “***broad clusters***” (**Figure 2d**). Note these clusters are agnostic to loci of origin, intended to clearly visualize differences in binding between HLA-A, HLA-B and HLA-C alleles. Indeed, upon closer examination of the ***broad clusters* (Figure 3 e,f**), a baseline level of promiscuity was observed throughout. The median recurrence rate of peptides between different regions of the map is 13%, with adjacent parts of the map having high rate of recurrent peptides, sometimes reaching 50% (**Figure 3f**).

The fact that peptides are promiscuous within a confined motif-space likely has an impact on models predicting antigen presentation – something CARMEN may help the machine learning community to grasp. In a general machine learning context, the “Hans effect” refers to the tendency of models to latch onto dominant, recurring patterns in the data, essentially memorizing these features, rather than fully learning the underlying complex relationships. We explored the adaptation of a deep learning model of antigen presentation, TransPHLA^27^, to scramble the HLA information on the datasets used to train/test TransPHLA (**Figure 3d**). We sequentially randomized the HLA or peptide columns in training, testing or both, as described in **Supplementary File 3,** to explore the reliance of the model on each parameter. The training and testing datasets consist of tuples of (peptide, HLA) with binding/not-binding response. While the model trained and tested on the unchanged TransPHLA source data is clearly better, randomization of the HLA in only training or both training and testing demonstrates AUCs > 0.9 to predict the same response, regardless of HLA Allele. Even randomization of just the test set HLAs, so still trained on correct data, still performs surprisingly well (AUC 0.7). We believe these surprising results are related to promiscuity, as there is some probability that the newly and randomly assigned HLA still binds the peptide by chance. It is interesting, that randomizing peptides destroys model performance (AUC ∼0.5), we believe randomizing HLAs still preserves a lot of information about promiscuous peptides. Randomizing peptides in context of the 3 components of the problem (HLA, peptide, response) completely ablates any information about promiscuous antigen binding. Hence, we believe, the observed performance may be attributed to the promiscuity of peptides, as reflected in their recurrence among the broad clusters (**Figure 3 e,f**). We hope the machine learning community will explore the consequences of promiscuity from a machine learning context more deeply, and that efforts will continue to understand the negative space of peptides that do not bind to specific MHC Alleles to improve models of antigen presentation.

### Promiscuity of MHC Alleles, individuals and populations

Overall, the low complexity of the revealed atlas has implications for similarities between individuals in antigen presentation. Considering the HLA motif atlas^24^ alleles depicted in **Figure 2d**, if every individual has 3-6 MHC Class I alleles representing motifs somewhere in this plot of physically identified peptides, the space may quickly saturate. The implication is an expected sample-to-sample similarity in antigen presentation between individuals and potentially populations. CARMEN clarifies the extent of this type of promiscuity as many samples in the CARMEN database have a known MHC Haplotype, and each sample creates several motifs that are associated with that sample’s haplotype.

#### Visualizing promiscuities and territorial biases of peptides connected to MHC alleles

We illustrate that the same definition of promiscuity can be applied to MHC Alleles. The MHC motif-atlas alleles^24^ are well approximated by the motifs of the CARMEN sample-derived Gibbs clusters of peptides present in our map, where territories in **Figure 2d** define motifs for different MHC Motif atlas^24^ alleles. Hence, we could approximate an MHC allele as a group of peptides with a shared motif at a coordinate on **Figure 2d**. These peptides aren’t exclusive to that allele/cluster — many are also found in the clusters corresponding to different MHC alleles. As a result, the entire group of peptides representing an MHC allele displays a high level of promiscuity, meaning it overlaps with multiple other clusters/motifs in the binding landscape. Indeed, the peptide clusters in our revealed map have a diversity of promiscuity scores indicating that peptides found associated with one motif pattern are also found widely in other Motif clusters in the map (**Figure 4a).** An extreme of promiscuity has been exemplified by the HLA C*12:03 allele from a cluster stemming from a mono-allelic cell line that binds peptides that are found throughout the landscape^28^. The promiscuity of each of the Gibbs clusters in the UMAP has been provided in **Supplementary Table 4**.

**Figure 4:**
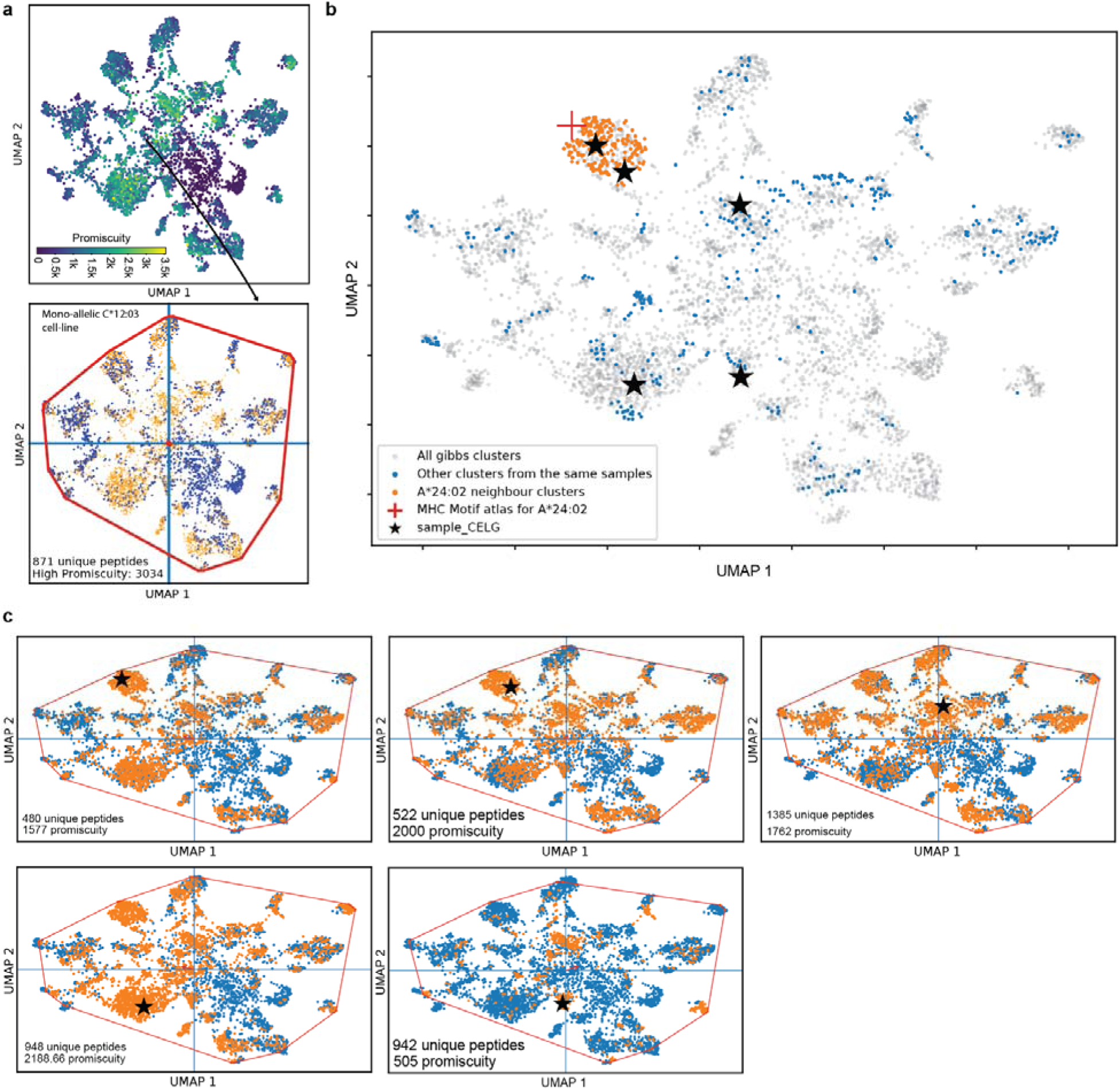
Promiscuity of HLA Alleles and Individuals. MHC Alleles bind onto a diversity of peptides, each of which has a promiscuity – occupying defined territories on the UMAP reflecting that each peptide binds also to other MHC Alleles. Hence the promiscuity of an MHC Allele, depends on the promiscuity of the peptides that it binds. **(a)** The heatmap illustrates the promiscuity of MHC alleles, represented by the diversity of Gibbs clusters that include some, but not necessarily all, peptides associated with the MHC allele within the landscape. The arrow illustrates a highly promiscuous Gibbs cluster in the map, containing peptides that map throughout the UMAP landscape. The cluster is derived from a mono-allelic cell line as indicated. **(b)** Each individual’s HLA alleles are visualized as constellations on the UMAP landscape, with one sample shown as an example, represented by a constellation of black stars. Up to six Gibbs clusters can form per sample, spreading across the landscape. The figure summarizes 68 samples with MHC Allele A*24:02, demonstrating that the samples reliably visit the appropriate UMAP neighbourhood for this allele, while their other alleles cause Gibbs clusters to appear in additional regions (blue). Building on the individual shown in (b), promiscuity is defined by the territories covered by each HLA allele in the sample **(c)**. The Gibbs clusters representing the sample exhibit a broad territorial reach, encompassing peptides identified within the sample and shared with clusters from other samples.

#### Understanding promiscuities and territorial biases of individuals as sets of MHC Alleles

CARMEN allows the visualization of individual immunopeptidomics samples as constellations of motifs on the revealed map (**Figure 4b)**. In theory, each HLA allele may yield a motif through a clustering of peptides in the sample by Gibbs Cluster and occupy some territory on the motif map. It is reassuring to see that nearly all samples sharing the same allele (A*24:02) visit the part of the UMAP with the correct MHC motif atlas^24^ allele (**Figure 4b**), but also interesting to see that these individuals comprehensively cover the motif space due to their diversity of haplotypes. The promiscuity of each Gibbs cluster emerging from a single sample in CARMEN has been visualized in **Figure 4c**, illustrating that the different motifs identified in an individual (**Figure 4b**) quickly raise the promiscuity to cover the landscape. Sample-wise promiscuity can be found in **Supplementary Table 5.**

#### Territorial biases in human populations revealed through sample promiscuities

MHC Class I allele frequencies are known to differ between populations based on the frequency of MHC haplotypes in each population. CARMEN can be used to understand biases in MHC Class I binding profiles of different human populations using our MHC Class I motif atlas. We explored both predictive approaches to comparing antigen presentation between populations to physical identification of peptides using the CARMEN dataset. Allele frequency distribution data from the NMDP Population Frequency database^29^, which contains information on MHC types for different populations in an organ donor registry was used in both cases. We used a table containing a total of 2,323 rows of MHC-I triplet allele frequencies across 21 different populations.

#### Approach 1: Using NetMHCpan 4.1 to reveal antigen-binding similarities within and across human populations

Intra- and inter-group immunopeptidome diversity was analyzed by sampling 3,000,000 random peptides (1,000,000 of lengths 8, 9, and 10 AA), with equal probability for each amino acid at each position. Peptide presentation by MHC-I alleles was predicted using NetMHCpan 4.1 (rank limit of 2%), allowing for the identification of peptides presented by an individual’s up to six MHC-I alleles by sampling two triplets of MHC Alleles according to their frequency in the individual’s population. Immunopeptidomes of two individuals generated in this way were compared 10,000 times for different pairs of populations, and unique and common peptides were identified (**Figure 5a**). On average roughly 35% of peptides were binders between individuals within and between any population, indicating that prediction perceives large common core of promiscuously presented peptides between individuals.

**Figure 5:**
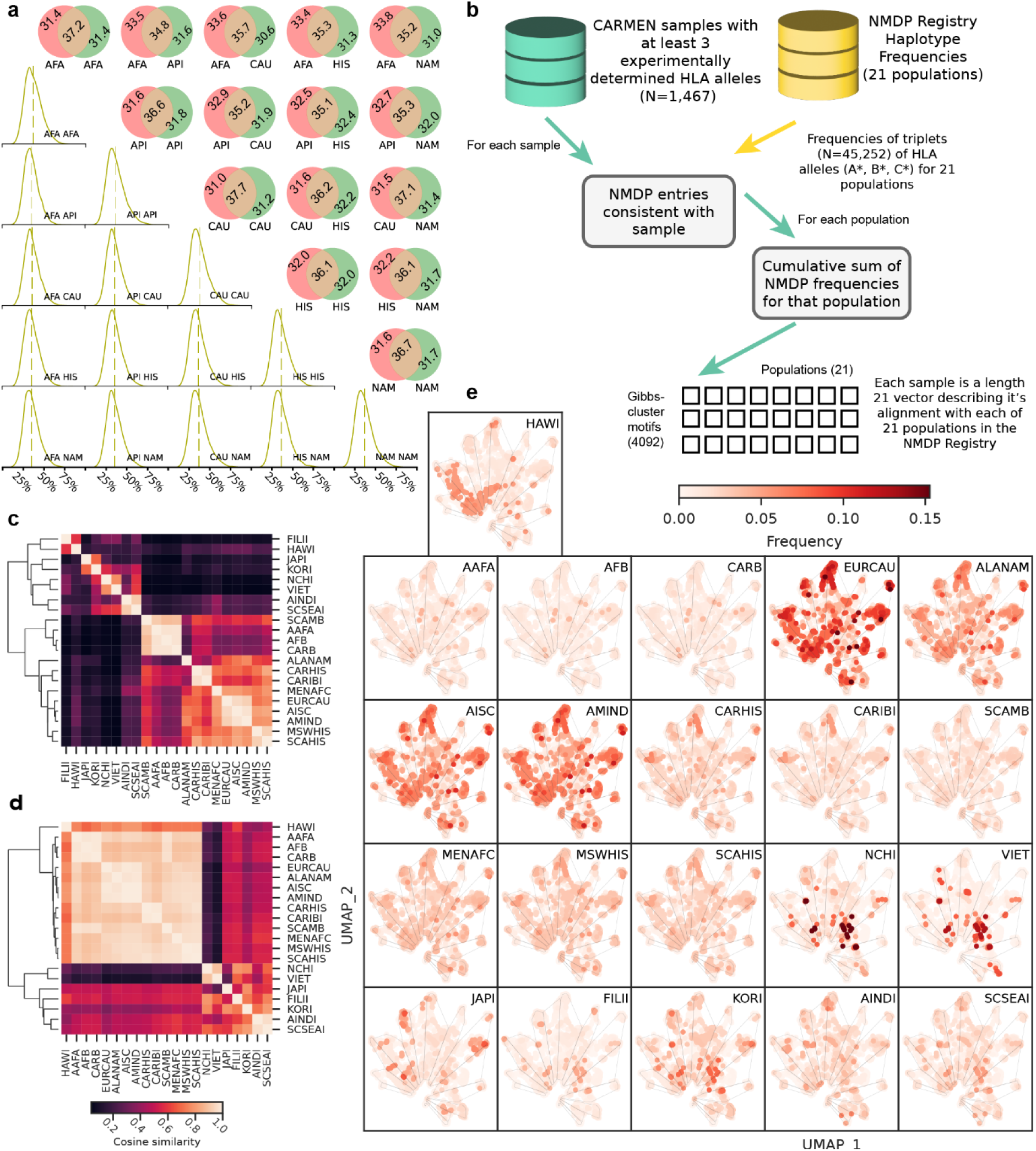
Promiscuity and territorial biases in human populations. We employed both predictive and physical detection approaches to compare peptide-binding biases across different population groups, accounting for the population-specific frequencies of MHC alleles. **(a)** We sampled 3,000,000 randomly generated peptides (1,000,000 of lengths 8, 9, and 10 AA). Using netMHCpan 4.1, the predicted presentation of these peptides by various MHC-I alleles (with a rank limit of 2%) was predicted. We used the NMDP Population Frequency database to simulate individuals with up to 6 MHC alleles from specific human populations. The Venn diagrams depict the average number of peptides shared between two individuals, while the histograms show the distribution of the proportion of peptides in the intersection. **(b)** The second approach determined how well aligned the allele frequency distribution data from the NMDP Population Frequency database (131) was between different human populations by first considering each populations alignment --likelihood to present similar peptides-- to motifs represented in the CARMEN UMAP landscape. Each population’s haplotype frequencies was aligned to the 4092 Gibbs clusters using the MHC allele information of the sample of origin to produce a 4092 motifs x 21 populations matrix -- where each entry represents the dot product to dot-product to the cumulative population frequencies. These long vectors were compared to each other, to produce a 21 x 21 matrix where each square of the heatmap represents the similarity in the biases of the UMAP motif coverage per population. **(c)** The overall population divergence as depicted by the haplotype frequencies database alone. and **(d)** with haplotype annotations within the CARMEN database. **(e)** Visualizing the territorial biases of different human populations at the sample level (each dot in an individual UMAP represents a motif and peptide cluster), showing the frequency of coverage in different populations. The more frequently the particular triplet-haplotype occurs in the population, the darker the colour.

#### Approach 2: Using experimentally verified binders from CARMEN

We measured the extent to which each sample in the CARMEN dataset was representative of each of the 21 populations in the NMDP Population Frequency database based on the samples with experimentally known haplotype (**Figure 5b**). For each sample with at least 3 MHC experimentally known alleles (n=1,467), the NMDP rows that fully matched the sample’s triplet-haplotype were aggregated to estimate the frequency at which this triplet-haplotype occurs within each of the 21 populations. We display the cosine similarity of the raw MHC Triplet allele frequencies in the NMDP database for the 21 different populations (**Figure 5c**) alongside the CARMEN observed sample alignment to each population (**Figure 5d**). In result, while some degree of divergence is observed across all populations, there are only two significantly distinct groups in terms of haplotype frequencies: populations native to the Asian continent (NICHI, VIET, JAPI, FILII, KORI, ANDI, SCSEAI) and all other populations. This distinction is particularly evident considering the haplotype data within the CARMEN database. Since each motif can be connected to a sample and its MHC haplotype, **Figure 5e** highlights each of the 21 calculated population frequencies calculated for each motif. The panel order here is consistent with the similarity clustering shown in **Figure 5d** and underlines the fact that most populations share similar antigen binding patterns as the same motifs are aligning to different populations, yet still some of them display differences. What is most interesting, is that we uncovered a significant bias in immunopeptidomics studies, as sample alignment was extremely high towards European Caucasians, highlighting a need to diversify the immunopeptidome (Supplementary Figure 2).

#### Promiscuity in genomic regions: Mapping peptides to contiguous regions reveals distinct clustering

We expand the concept of promiscuity to encompass genomic regions by mapping the immunopeptidome to the genome. We collapsed mapped peptides into contiguous “epitope contigs” and higher-level “epitope scaffolds” using variable gap lengths (10–300 nucleotides; 3 to ∼100 amino acids) (**Figure 6a**, **6b**). Peptides were directly linked to genomic coordinates for seamless integration with genomic tools, with contigs and scaffolds available in BED format.

**Figure 6:**
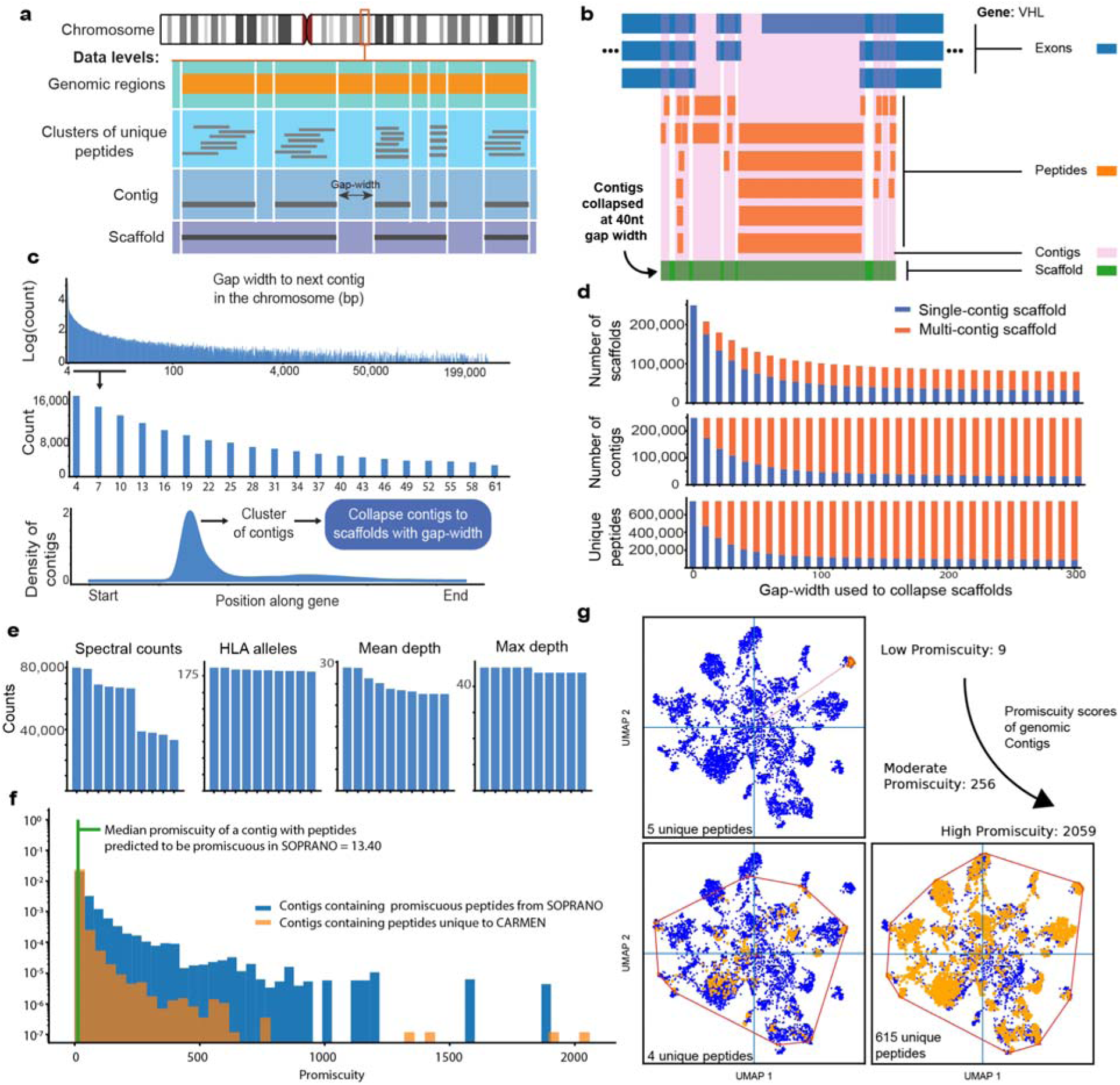
Promiscuity of genomic regions. Mapping peptides to the genome allowed us to better understand the promiscuity of genomic regions. **(a)** After mapping peptides to the genome, we developed contigs composed of contiguous clusters of these peptides and collapsed contigs to scaffolds at a pre-specified gap-width. **(b)** The VHL gene (ENSG00000134086) is used to illustrate the collapsing of the different levels of the genomic model and visualise the levels at each instance. **(c)** The distribution of the log-scaled number of contigs per gap width (top), with the most abundant regions zoomed in (middle) and an example gene (ENSG00000182901) with clusters and start/end coordinates described (bottom). **(d)** The frequency distribution of different types of scaffolds per gap-width is visualised together with the number of contigs and unique peptides seen in each contig. **(e)** The scaffolds with the maximum values of spectral counts, HLA alleles, mean and max depth. **(f)** To define a contig-level promiscuity cut-off, we integrated CARMEN-derived contigs with SOPRANO-identified regions^26^ based on peptides predicted to bind the top 75 most frequent MHC alleles. **(g)** Examples of highly and lowly promiscuous genomic regions with their promiscuity scores and number of unique peptides physically detected in these regions. Each dot on the UMAP represents a sample within the larger Gibb’s cluster motif space.

There was a bias to find contigs near each other on the genome. Analyzing peptide clustering, we found a bias toward shorter gap lengths (0–20 nucleotides, ∼8 amino acids) (**Figure 6c**). Smaller gap widths collapsed more scaffolds while increasing unique peptides (**Figure 6d**). Epitope scaffolds assembled at a 40-nucleotide (∼13 aa) gap covered 10% of exons in most genes (47, 870), increasing logarithmically from 16.84% to 41.18% with gap widths up to 300 nucleotides. Of 248,526 contigs, 51.1% clustered within 61 nucleotides (∼20 aa), 11.8% within 10 nucleotides (∼3 aa), with multi-contig scaffolds peaking at 50 nucleotides (∼16 aa) (**Figure 6d**).

We present the top 10 scaffolds at 50 nt (∼16 aa) (**Figure 6e**). Notably, regions exposed to the immune system often had nearby exposed regions within the CARMEN dataset. Contigs exhibited varying promiscuity across the dataset, exemplified in **Figure 6f**. To define a contig-level promiscuity cut-off, we integrated CARMEN-derived contigs with SOPRANO-identified regions^26^ and quantified their overlap (**Figure 6f**). We set at the promiscuity level of 13 below which overlap to SOPRANO declined and designate promiscuities larger than this value as *promiscuous contigs*. We also highlight three types of contigs with varying promiscuities in **Figure 6g** showing that as more unique peptides cover a genomic region, the MHC allele territory and therefore promiscuity of that region also increases.

### Cancer Mutation Landscapes and Immunotherapy Response

We next examined how peptides, contigs and scaffolds could help understand and predict immunotherapy response by integrating COSMIC mutation rates as well as contig features like unique peptide abundance, and promiscuity, bridging immunopeptidomics with cancer mutation data.

### Mutations, Promiscuity and Immune-Visibility

We ranked the top 50 recurrent non-synonymous mutations from the COSMIC Cancer Gene Census by contig promiscuity, focusing on promiscuous regions that ranked above the median promiscuity and unique peptide abundance (**Figure 7a**). Notably, highly recurrent mutations often occur in low-promiscuity contigs, making them less ideal for cancer vaccines. The contigs harbouring the most promiscuous recurrent mutations, IL32-D172Gfs and TP53-H193R, occupy distinct immune landscapes (**Figure 7b**). Several highly promiscuous TP53 regions and hotspots in CTNNB1, TP53, and KRAS also emerge in this ranked list.

**Figure 7:**
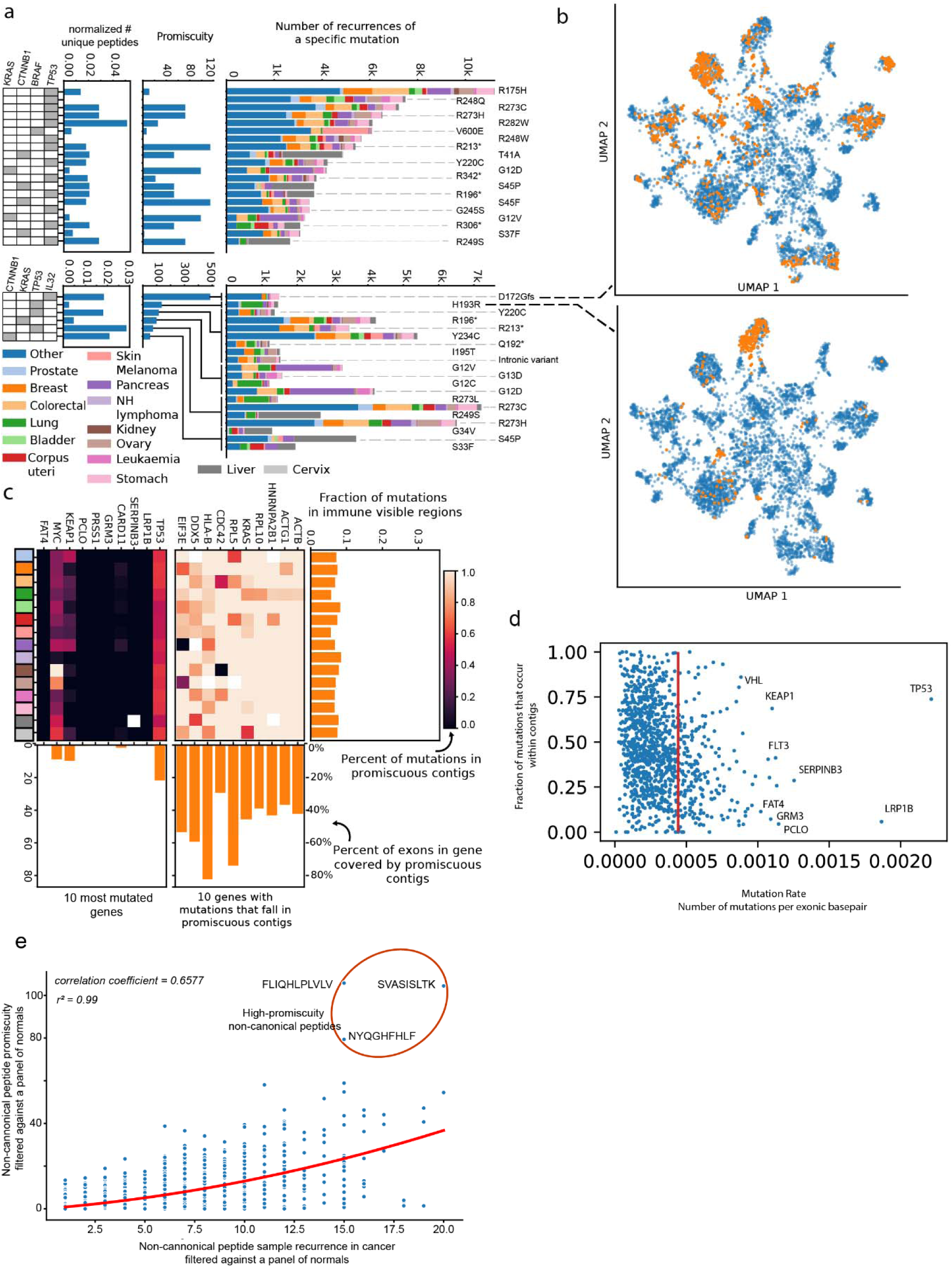
Promiscuity and recurrence in mutations and genes in cancer. **(a)** The number of recurrent mutations observed in COSMIC within each contig region and their associated features (promiscuity & unique peptides) for all regions (top) and those in the top quadrant of contigs defined as contigs with promiscuity greater than 13 (highly promiscuous) and number of unique peptides above the median number (bottom). For context, the pathological grouping of the COSMIC samples containing these mutations and the genes within which they are found is displayed. On the right **(b)** are displayed UMAPs for two representative mutations falling in highly promiscuous contigs falling in IL32 (top) and TP53 (bottom); each blue dot represents a Gibb’s cluster motif and each orange dot represents clusters containing peptides forming the contig overlapping with given mutation. **(c)** Some cancer related genes are more immune-visible than others. Immune-visibility of the most mutated genes in cancer (left heatmap) and the genes with the greatest number of mutations found within promiscuous immune visible contigs. The analysis was performed across COSMIC database samples matching the 15 highest incidence cancer types according to ECIS (European Cancer Information System) data. **(d)** Mutations in important cancer genes defined by (oncoKB) fall along a spectrum of immune-visibility. We annotate the fraction of mutations that fall within contigs in a gene and the exonic region mutation rate, with genes showing the highest number of both exonic mutations and per-contig mutational fraction highlighted. The red line represents the top 20’th percentile of genes by their exonic region mutation rate.

Certain highly mutated genes are more immune-visible than others (**Figure 7c**). While mutations in certain genes are often immune-visible, many highly mutated cancer genes (OncoKB^30,31^) have mutations that fall outside contigs. Genes such as ACTB, ACTG1 HNRNPA2B1, RPL10, KRAS, RPL5, CDC42, HLAB, DDX5 and EIF3E are among those that exhibit greater immune-visibility, suggesting their relevance as targets for immunotherapy and cancer vaccines (**Figures 7d**), but even in these just 10% of mutations fall within the promiscuous contigs.

### Non-Canonical Immunopeptidome Promiscuity

Building on Bedran *et al.* (2023)^20^, we refined the non-canonical immunopeptidome landscape in CARMEN with a more comprehensive normal panel using the HLA-ligand atlas^32^ which is among the datasets reprocessed in this work. Non-canonical peptides exhibit diverse promiscuities (**Figure 7e; Supplementary Table 8**), with SVASISLTK as a highly promiscuous example.

#### Promiscuous contigs in predicting response to immunotherapy

To assess the impact of promiscuous contigs on immunotherapy response, we trained a Support Vector Machine (SVM) classifier using attributes including unique peptides, HLA alleles, spectral counts, expression depth, and promiscuity. Our analysis revealed diverse immune visibility (promiscuity) across genomic regions differently associated with different rates of cancer-specific mutation as observed in COSMIC (**Figure 8a**). We then evaluated whether mutations overlapping with previously detected immune-peptides, contigs or scaffolds predict immunotherapy response at the population level. An SVM classifier incorporating peptides, contigs, and scaffolds predicted Progression-Free Survival (PFS) in 57 ccRCC patients treated with Nivolumab, distinguishing responders (PFS ≥ 6 months) from non-responders (PFS < 6 months) (**Figure 8b**). Three kernel functions—linear, radial basis function, and polynomial (degree 2)—were tested. Various feature sets and weightings, including prognostic scores (PS), biological processes (BP), tumor/clinical features (TF/CF), mutations (MT), and gene expression (GE), were evaluated (see **Supplementary File 3**).

**Figure 8:**
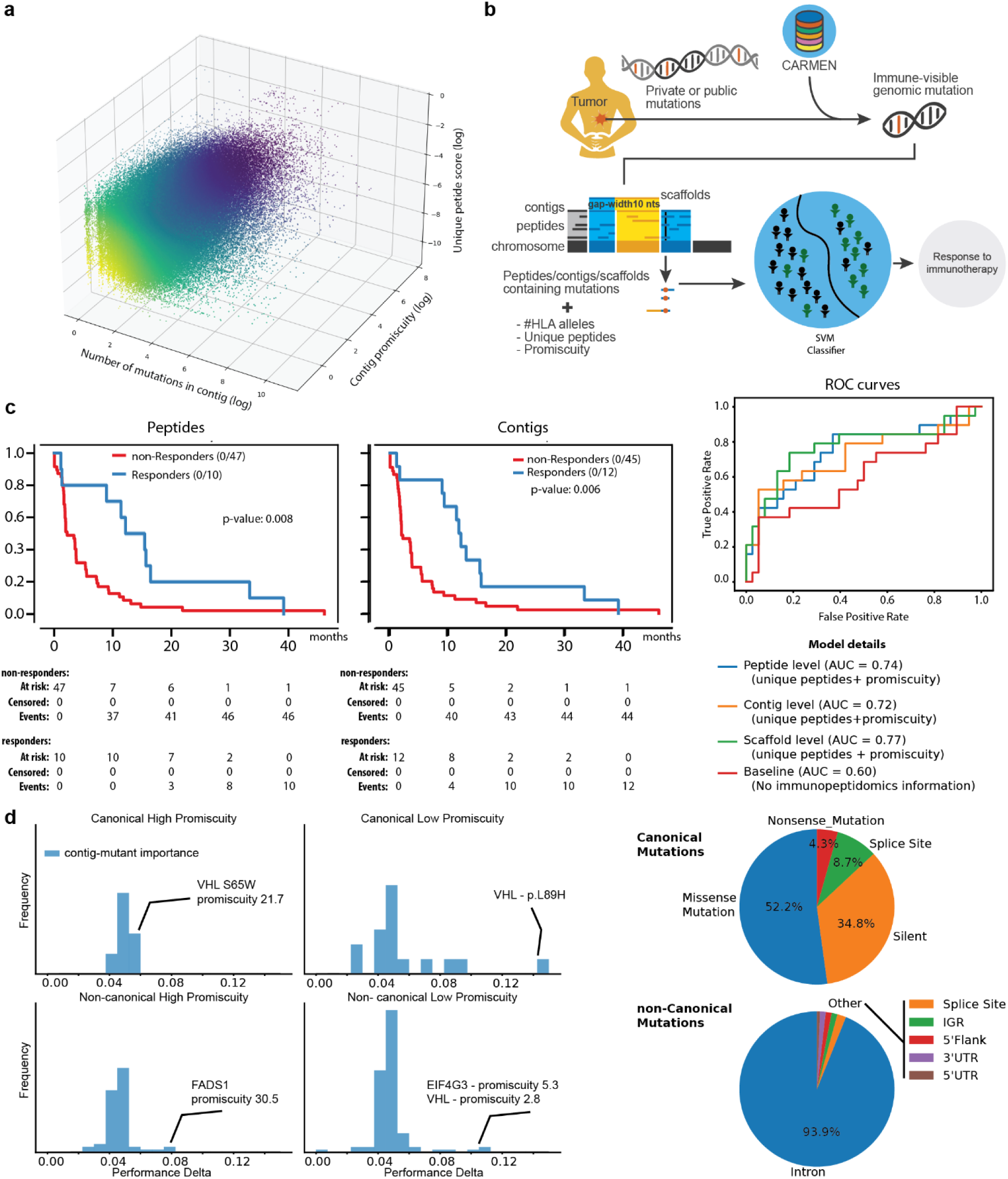
Promiscuity and implications for immunotherapy and cancer vaccine design. **(a)** Distribution among contigs of three features tied to each contig: promiscuity scores, mutation counts and unique peptides. **(b)** An SVM classifier was developed using patient data obtained from the Braun cohort (2020), comprising 311 clear cell renal cell cancer (ccRCC) patients who had previously been treated with nivolumab. We explored the value added of the three features in (a) tied to peptides overlapping the mutation, contigs and scaffold-level data in predicting immunotherapy response. **(c)** Predicted overall patient survival for well performing models built considering peptides directly overlapping patient mutations, as well as more generally the contigs that they contain are shown. ROC curves including the scaffold level results show better performing ROCs over base-line models that do not include immunopeptidomics data. **(d)** The importance of mutations within the contig model in predicting immunotherapy response by computing changes in model AUC performance. The performance delta AUC score for canonical and non-canonical mutations within promiscuous and non-promiscuous contigs is shown.

The classifier stratified patients with varying accuracy based on promiscuity level, attribute selection, and weighting. In all cases mutations recurrent in at least 4 patients were used. Peptide-level classification was significant (log-rank p = 0.008), as were the contigs (p = 0.006) worsening for scaffold levels (p = 0.02) (95% CI) (**Figure 8c**). ROC statistics calculated on 6-month progression free survival indicate that adding information about genomic regions of interest always produced improvements over models that did not, particularly the number of unique peptides and promiscuity of the region. The AUROC improved from 0.60 (baseline p=0.02) to 0.77. Weighting biological processes, mutations, and clinical features further enhanced AUROC and log-rank scores. Compared to the original nivolumab-treated cohort’s PFS baseline (**Extended Data 8d from Braun *et al.***), our multi-parametric SVM model showed improved predictive power, and also compared to our own baseline, ignoring the immunopeptidome, which has a p-value of 0.02. Given the similar predictive performance of peptides, contigs and scaffolds, key response-associated contigs were identified by analysing AUROC drop (**Figure 8d**). We explored the promiscuity of contigs harboring important canonical and non-canonical mutations – and suggest this information can be used to better understand response to immunotherapy. For example, the VHL gene, critical in ccRCC progression^33,34^, ranked among the top ten most predictive contigs, but in a diversity of promiscuities. Full results are in **Supplementary Table 7**.

## Discussion

Promiscuity in T-cell antigen landscapes refers to MHC Class I and II molecules’ flexibility to bind various peptides and the ability of certain peptides to be presented by multiple MHC alleles. This concept could help to interpret immunotherapy responses by mapping the binding territories of neoantigens from recurrent mutations, highly mutated genes, and tumor-associated peptides. Such an understanding may help in prioritizing precision cancer vaccines and evaluating neoantigens for off-the-shelf vaccines, potentially enabling broader population coverage and streamlined treatments^35^.

CARMEN (**Figures 1,2**) expands the promiscuity concept (**Figures 1h, 2de**) for broader applications, providing comprehensive data to compare binding profiles of peptides (**Figures 2,3**), MHC alleles (**Figure 4a**), individuals (**Figure 4b,c**), populations (**Figure 5**), and genomic regions (**Figure 6**). While the MHC-I motif landscape is considered complex^24^, studies^13–15^ suggest significant promiscuity. We clarify this by mapping MHC Class I motifs across 2,323 samples (**Figure 2d**), tracking peptide membership to clusters in the revealed map. Reassuringly, our findings demonstrate that samples with identical MHC alleles cluster in specific regions of this map (**Figure 4b**).

### CARMEN’s motif landscape can be used to understand promiscuity in different contexts

The motif landscape of physically detected MHC Class I peptides is surprisingly simple (**Figure 2d**). While our samples show wider motif variation, the overall motif space appears to be saturating. Peptides exhibit diverse promiscuities (**Figure 3**), and the motif landscape quickly saturates when considering MHC allele and individual promiscuity (**Figure 4**). Since patients may have up to 6 MHC alleles representing motifs in our dataset (shown in **Figure 4b**), peptide binding promiscuity between individuals is amplified.

Predictive modeling demonstrates this amplification, estimating that over one-third of peptides are shared between any two individuals across all populations (**Figure 5a**). While different populations emphasize certain aspects of the map, significant overlaps exist among most populations, with East Asians showing clear distinction (**Figure 5e**). Although HLA allele distribution varies globally^36,37^, with documented locus-specific variations among East and Southeast Asian populations^38–40^, sample haplotypes in current datasets show extreme bias toward European Caucasian populations, highlighting the need for greater diversity in immunopeptidomics studies.

Recognizing promiscuity at individual and population levels is crucial as neoantigen selection relies heavily on predictive modeling. These models can both over- and underestimate peptide presentation, with newer transformer-based methods potentially prone to memorization due to limited negative binding data. Notably, models (not based on our data) that scramble MHC allele data still perform very well (**Figure 3d**). CARMEN provides the genomics community with comprehensive data on physically detected peptides and their promiscuity across mass-spectrometry studies, complementing predictive methods. Promiscuous immunopeptides with high population coverage and immune visibility are valuable for response prediction and vaccine design, offering potential for broader population application.

### Promiscuity of genomic regions

We examined “hotspots of antigen presentation” through promiscuity and antigen binding “territories.” Genomic regions can be defined by their mapping MHC binding peptides, with promiscuity visualized by motifs found in our landscape (**Figure 6**). When peptides map to the genome, the next mapped peptide, often from a different sample, frequently appears nearby (**Figure 6 c,d**). This observation led to our definition of “epitope contigs” (contiguous regions mapped by MHC Class I peptides) and “epitope scaffolds” (semi-contiguous regions where peptides may be separated by “gap-lengths”). We explored various scaffold gap lengths (**Figure 6d**) and include these definitions in the dataset accompanying this manuscript. These regions exhibit diverse promiscuities (**Figure 6g**) based on their unique peptides, HLA allele membership, and extent to which they are mutated in cancer (**Figure 8a**).

### Some cancer genes are more immune-visible than others

The fact that genomic regions are differently promiscuous highlights differential surveillance across individuals (per CARMEN samples) and suggests population genetics may self-regulate regional promiscuity. Cancer genes can be classified by whether mutations occur within contigs. Tumor suppressors TP53, KEAP1, and VHL have exons covered by 25%, 26%, 28%, and a large fraction of mutations in these genes appear within contigs (**Figure 7d**). Conversely, often mutated cancer genes LRPB1B, FAT4, GRM3, and PCLO rarely have mutations overlapping regions that appear in the immunopeptidome. The most “immune-visible” genes with highest exonic coverage in promiscuous contigs include ACTB, ACTG1, HNRNPA2B1, RPL10, KRAS, RPL5, CDC42, HLA-B, DDX5 and EIF3E (**Figure 7c**). With >40% exon sequence coverage in CARMEN, these genes may be ideal candidates for gene-specific poly-epitope vaccines.

### Promiscuity of recurrently identified mutations and non-canonical peptides in cancer

The immunopeptidome’s simplicity supports the public neoantigen^35^ hypothesis, particularly given landscape saturation across individuals’ six MHC alleles (**Figure 4**) and demographic similarities (**Figure 5**). We’ve released promiscuity data for contigs containing recurrent cancer mutations, finding several (IL32, TP53, KRAS) within promiscuous contigs (**Figure 7a**). However, most recurrent mutations show variable promiscuity, suggesting reduced immune visibility of public neoantigens. In addition, CARMEN extends our previously revealed non-canonical peptide landscape^20^, filtered against an improved normal database^32^, with promiscuity annotations shown in **Figure 7e**. We caution that we do not apply gene-expression filters as in the original publication, as our non-canonical peptides are derived from non-coding regions of existing transcripts – and we believe there are better experimental methods that could be applied to assess toxicity.

### Population-level trends in response to immunotherapy can be predicted by certain contigs

Our analyses revealed that epitope contigs can aid the prediction of immunotherapy response and we observe that mutations in certain contigs assume greater importance than others (**Figure 8**). We were interested in understanding intronic peptide presentation, so we designed our database such that the intron was retained in contigs with peptides overlapping a splice-site. For this reason, we were able to explore the promiscuity of both coding and intronic mutations in **Figure 8d**. We highlight the most important canonical and non-canonical mutations in different promiscuity contexts (**Figure 8d**), which may add some explainability to VHL mutations of import to renal cancer.

### Contribution of CARMEN to the landscape of immunogenomic databases

CARMEN contains 816,222 unique peptides from 31 cancers and 2,323 samples, offering a proteogenomic perspective distinct from databases like caAtlas^41^, SysteMHC Atlas^42,43^, SYFPEITHI^44^, Tantigen^45^, IEDB^46^, CEDAR^47^, and HLA Ligand Atlas^32^. While IEDB and CEDAR aggregate multi-species data across B/T-cell assays, CARMEN focuses on promiscuity in human MHC Class I antigen binding. Unlike cancer-centric CEDAR, healthy-tissue-exclusive HLA Ligand Atlas, or multi-disease IEDB/SysteMHC Atlas, CARMEN enables comparative analysis across cancer, healthy, and benign tissues. It includes 8,500 non-canonical peptides (from non-standard reading frames) and integrates with FAIR-compliant Immunopeptidomics Ontology (ImPO)^48^. CARMEN clarifies MHC Class I allele motif usage, revealing 90% of immunopeptide space is covered by just 3.35% of possible amino acid pairs at specific positions (**Figure 2a**), with leucines dominating positions 2 and 9. This high-resolution motif analysis extends to 8-amino-acid combinations (**Supplementary Figure 1**), providing unprecedented insight into protein exposure mechanisms.

### Concluding remarks

Our work redefines and clarifies “promiscuity” to include haplotypes, peptides, and genomic regions, demonstrating its utility in predicting immunotherapy response and immune visibility. These regions often overlap immune-visible mutations and may be differentially targeted by the immune system. Supporting this, a study^49^ reported improved survival in nine ccRCC patients treated with vaccines targeting VHL, PBRM1, and PI3K mutations - mutations aligning with our highly promiscuous peptides. Future research should validate these findings using a multi-omic approach to assess their impact on patient survival. Our proteogenomic model could transform immunopeptide identification, vaccine design, and therapy prediction by integrating multi-omic factors, enabling both patient-specific and broad-coverage treatments. We aim to pioneer this integration, setting our approach apart in the field.

## Data availability

The mass spectrometric datasets reanalysed are freely available on PRIDE and MASSIVE and are listed in **Supplementary Table 1**. Please note that while our intention is to release the data as openly as possible, this dataset is a compilation of data from multiple sources. Some source data is shared under licenses that restrict commercial use or have other conditions. As a result, this compiled dataset is not licensed under a single open license. The end user is responsible for reviewing and complying with the individual licenses of all source data before using this dataset, especially for any commercial, redistributed, or derivative use. A list of source datasets has been provided in the supplementary tables. The code used for data analysis, engineering, and visualization has been deposited at the following public GitHub repository: https://github.com/immuno-informatics/carmen-analysis. All supplementary tables, figures, and files are available at Zenodo: https://doi.org/10.5281/zenodo.14859003, where a zip file entitled “Supplementary Materials Manuscript.zip” can be found which conforms to the tables referred to in the manuscript. The CARMEN database tables have been deposited to Zenodo under the following link: https://doi.org/10.5281/zenodo.13928441. Because of the dedication to open-research, the published article is licensed under Creative Commons Attribution 4.0 International.

## Acknowledgements

We gratefully acknowledge the collective contributions that made this work possible. The co-first authors of this manuscript AAK, MW, AP and EDW contributed equally to this work and can exchange authorships on their CVs to acknowledge this – with appropriate credit given below. AAK, MW and AP reprocessed raw mass spectrometry data from PRIDE and MASSIVE using a previously established codebase^31^, while JA, CP, AP, MW and AAK curated the CARMEN database. EDW and AP led the motif analysis and Gibbs clustering and UMAP visualization. MW developed the promiscuity scoring method and performed the analysis for peptide-, HLA allele-level and individual sample promiscuities. AP modelled peptide presentation using haplotype frequency data to reveal population biases. FMZ, MM, DV, PB and AP compared predictive binders with experimental observations. Genomic mapping, including the design of epitope contigs and scaffolds, was developed by AAK, AP, MW, with statistic figures provided by AAK and MW. MW integrated COSMIC mutation data with CARMEN-derived contigs, with clinical input from MS, SS, TH, KP and AL. FMZ, PB, and MM developed the ML classifier to performed survival analysis. MW and AAK applied the SVM model to examine how mutations in promiscuous regions associate with immunotherapy outcomes, MW performed the parameter optimization. The conceptual framework and expanded definition of promiscuity in immunopeptidomics were defined by JAA and MW. Manuscript drafting by AAK, MK, MW, AP, EDW and JAA. Multidisciplinary integration was managed by JAA, AAK, MW and FMZ. We also acknowledge the significant consortium-wide contributions of The KATY Consortium. Some aspects of this work made use of GNUParallel^50^.

## Funding Acknowledgements

This work is supported by the Knowledge At the Tip of Your Fingers: Clinical Knowledge for Humanity (KATY) project funded from the European Union’s Horizon 2020 research and innovation program under grant agreement No. 101017453. We gratefully acknowledge Polish high-performance computing infrastructure PLGrid (HPC Center: ACK Cyfronet AGH) for providing computer facilities and support within computational grant no. PLG/2023/016653. AL is supported with funding from an MRC Clinical Academic Research Partnership Award (grant number MR/W030322/1).

## Methods

### Promiscuity in genomic regions: Describing the CARMEN dataset

We mapped peptides to the genome **Figure 5a**. We developed a proteogenomic data model linking peptide sequences to their genomic coordinates, enabling the integration of antigen presentation statistics with common genomic tools. The model consists of four views: Layer 1 maps peptides to genomic regions, tracking unique peptides and peptide spectrum matches (PSMs). Layer 2 provides protein-level hotspots, adding protein coordinates for each peptide. Layer 3 collapses peptide clusters into “epitope contigs,” representing continuous stretches of peptides at specific genomic locations. Layer 4 groups contigs into “epitope scaffolds,” collapsing using different gap lengths up to 300 nucleotides for higher-level organization. The model maps genomic regions across 2,323 samples, capturing key statistics like frequency of observation. It was visualised using the VHL (ENSG00000134086) (**Figure 5b**) and RGS7 (ENSG00000182901) genes (**Figure 5c**).

The following statistics were associated with contigs and scaffolds (**Figure 5d-e**):

a. *Unique peptides*: The number of unique peptides that fall into the region.
b. *Unique HLA alleles*: The number of unique HLA alleles presenting each of these peptides.
c. *Sum of normalized spectral counts*: The quantile normalized sum of all unique spectra mapping to each peptide within a contig.
d. *Mean & Maximum depth*: Referred to as peptide-level depth, this is the average and maximum per-position coverage of unique peptides across the contig.
e. *Width*: The difference between the starting and ending coordinates of the contig.

### Cancer mutation landscape through the lens of contigs and promiscuity

The recurrent mutations across the cancers described in COSMIC were tabulated and ranked based on (a) promiscuity score and (b) normalized unique peptides (**Figure 7a**, **7b**). The fraction of mutations within immune-visible regions was computed, followed by a correlation analysis between the percentage of mutations in contigs and the total number of mutations per gene (**Figure 7c**). Subsequently, a ranked list of genes by (a) most “visible” mutations, (b) least “visible” mutations, and (c) most mutations was tabulated, accompanied by a calculation of the percentage of exonic regions within contigs for each of these genes (**Figure 7d**, **7e**). The analyses were performed those most promiscuous mutations. Lastly, the promiscuity scores of two genes, namely (a) KRAS and (b) TP53, were visualized across all samples (**Figure 7f**).

To find the peptides presented from mutated regions in the studied cohort first the mutation data was translated to hg38 coordinates using liftOver^51^, and subsequently, a join was performed to find the intersection between the mutation regions and epitope scaffolds in the database, as well as between mutation regions and known presented peptides in the database.

To assess an individual patient, the peptides overlapping mutated regions were analysed against the motif space in the database, and a list of motifs they are a part of was obtained, which can be further narrowed down if the patient’s HLA haplotype is known.

The same scoring formula can be used to assess the immune visibility for a gene, in which case the peptide list to be intersected with the motif space is created by all the peptides from the database occurring within the gene, as well as an individual epitope contig, for which the peptide list is just the list of peptides forming the epitope scaffold. Using TP53 as an example, distinct clusters of immune visibility could be observed at contig regions across the gene as revealed by UMAP visualizations. The recurrence of non-canonical peptides was found to moderately correlate with their promiscuity (correlation coefficient 0.6575 using a polynomial regression fit, r-squared value: 0.99). The moderate correlation between recurrence and peptide-level promiscuity, even after excluding normal tissues, could potentially be explained by the disproportionate prevalence of peptides in some cancers (such as melanoma, colorectal cancer and glioblastoma) and the significantly lower prevalence in others (such as clear cell renal cell carcinoma). Despite this, certain peptides with high promiscuity and recurrence could still be identified (**Figure 7e**) (**Figure 2e**) which were interesting since they were also exclusive to cancer.

### SVM Classifier and survival analysis

A Support Vector Machine (SVM) model was developed incorporating region features (number of unique peptides, number of unique HLA alleles, Promiscuity) and tested to predict Progression Free Survival (PFS) in the context of immunotherapy based on the nivolumab arm of the Braun cohort (**Figure 8b**)^19^. We explored calculating features, including promiscuity, about immunopeptidomics data in different ways. Mutations in that dataset were enriched with information describing their epitope regions. Four levels of specificity were evaluated – In the ***peptides*** model, CARMEN peptides directly overlapping the mutation were used to calculate region features. In the ***contigs*** model, metrics for the epitope contig were used. Finally, in ***scaffolds models*** the scaffolds at gap widths of 10 and 20 respectively were tested. Each specificity level was evaluated for at least 10,000 random combinations of patient data scaled by random weights and with or without combinations of additional mutation information. As demonstrated the additional features provided for mutations within epitope regions demonstrated superior predictive power for immunotherapy outcomes compared to unenriched mutation information, independent of the kernel used. Survival analysis indicated that combination of unique peptides, and promiscuity (**Figure 8c**) outperformed individual features in predicting patient survival post anti-PD1 therapy for each of the studied specificity levels. The importance of each mutation and its contig meta-data in predicting immunotherapy response was determined using a performance delta on the AUC, with the greatest drop in performance indicating a significant contribution to immunotherapy response prediction (**Figure 8d**). This layers potential explainability to the importance of mutations in the context of response to immunotherapy in renal therapy, though we note that these correlations may not be causal. More detailed methods can be found in **Supplementary File 3**.

## List of abbreviations

MS: Mass Spectrometry
MHC: Major Histocompatibility Complex
HLA: Human Leukocyte Antigen
CARMEN: Cancer immunopeptidogenomics
UMAP: Uniform Manifold Approximation and Projection
PRIDE: Proteomics Identification Database
MASSIVE: Mass Spectrometry Interactive Virtual Environment
DDA: Data Dependent Accquisition
COD-Dipp: Closed-Open-de Novo-Immunopeptidomics
BED: Browser Extensible Data
F.A.I.R: Findable Accessible Interoperable Retrievable
PSSM: Position Specific Scoring Matrix
BLOSUM: Block Substitution Matrix
KLD: Kullback-Leibler Distance
SOPRANO: Selection in Protein Annotated regions
AUC: Area under the curve
ROC: Reciever Operating Characteristic
AUROC: Area under the receiver-operator characteristic curve
NMDP: National Marrow Donor Program
TransPHLA: Transformer Peptide HLA
COSMIC: Catalogue Of Somatic Mutations in Cancer
PFS: Progression Free Survival
PS: Prognostic Score
BP: Biological Processes
CF: Clinical Features
TF: Tumour Features
GE: Gene Expression
MT: Mutations
CI: Confidence Interval
caAtlas: Cancer Antigen Atlas
IEDB: Immune Epitope Database
CEDAR: Cancer Epitope Database And Analysis Resource
ImPO: Immunopeptidomics Ontology
ccRCC: Clear Cell Renal Cell Carcinoma
PSM: Peptide Spectrum Match

## List of supplementary tables

| S.No. | Table name | Table description |
| --- | --- | --- |
| 1 | Supplementary Table 1 | List of MS/MS datasets used for analysis |
| 2 | Supplementary Table 2 | Peptide counts per sample |
| 3 | Supplementary Table 3 | Experimental MHC haplotypes per sample (where known) |
| 4 | Supplementary Table 4 | Promiscuity of each Gibb's cluster |
| 5 | Supplementary Table 5 | Promiscuity of each sample |
| 6 | Supplementary Table 6 | Promiscuity of each peptide |
| 7 | Supplementary Table 7 | Performance delta values for all patients |
| 8 | Supplementary Table 8 | Promiscuity of non-cannonical peptides |

## List of supplementary figures

| S.No. | Figure name | Figure description |
| --- | --- | --- |
| 1 | Supplementary Figure 1 | All amino acid combinations at various positions |
| 2 | Supplementary Figure 2 | Frequency distributions of population HLA allele frequencies |

## List of supplementary files

| S.No. | File name | File description |
| --- | --- | --- |
| 1 | Supplementary File 1 | Closed search parameter file for MSGF+ |
| 2 | Supplementary File 2 | Open search parameter file for MSFragger |
| 3 | Supplementary File 3 | Supplementary methods |

