## Supplementary material for "Expanding the definition of MHC Class I peptide binding promiscuity to support vaccine discovery across cancers with CARMEN": Supplementary Figure 1.pdf

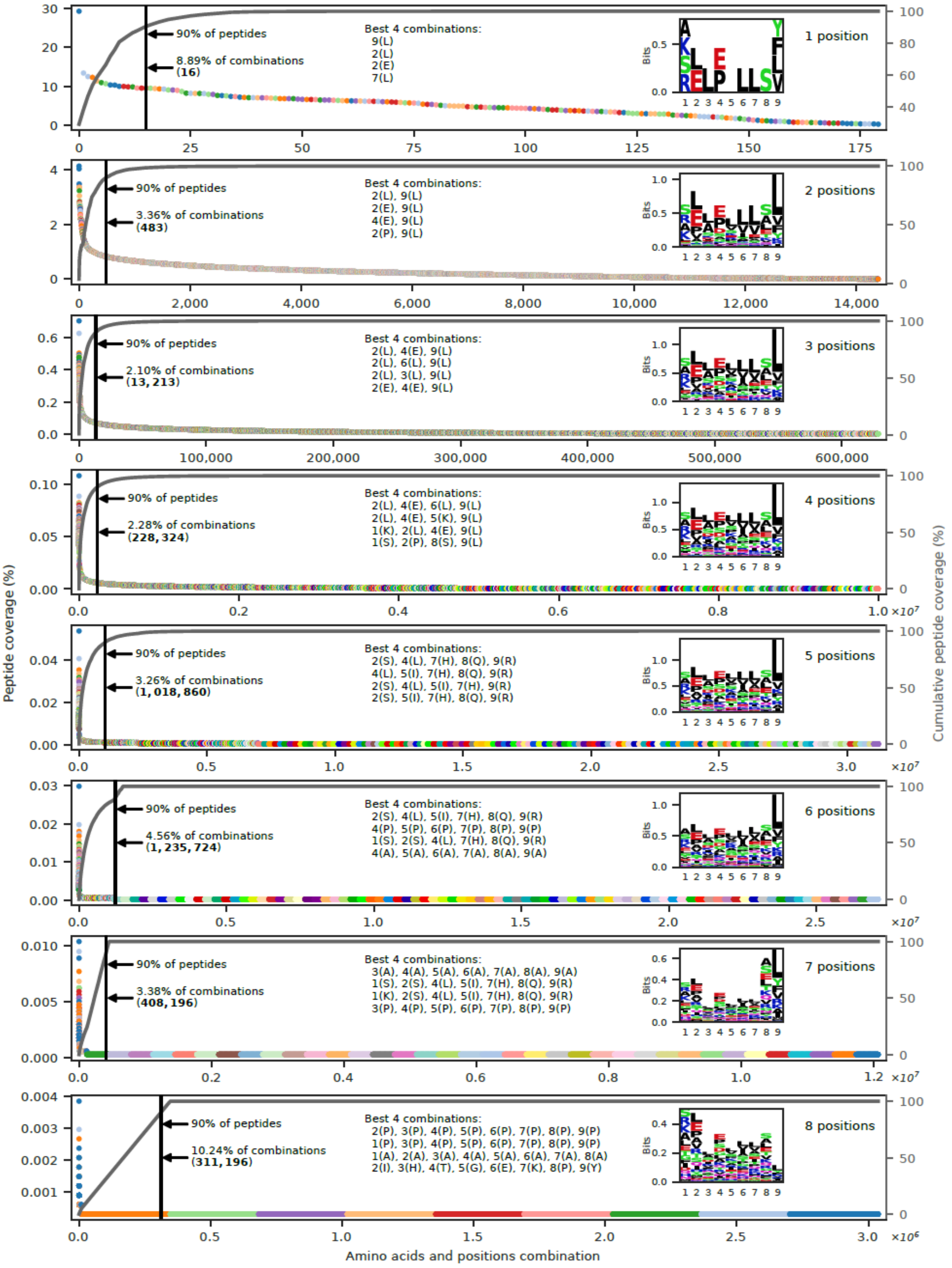


**Supplementary Figure 1:** The frequency of various amino acid combinations at specific positions within the peptide, showing the dominant motifs/residues per position per combination. We continued this analysis to combinations of 8 amino acids, obtaining a restricted space of fixed amino-acid position frequencies important for peptide presentation.
