## Supplementary material for "Expanding the definition of MHC Class I peptide binding promiscuity to support vaccine discovery across cancers with CARMEN": Supplementary Figure 2.pdf

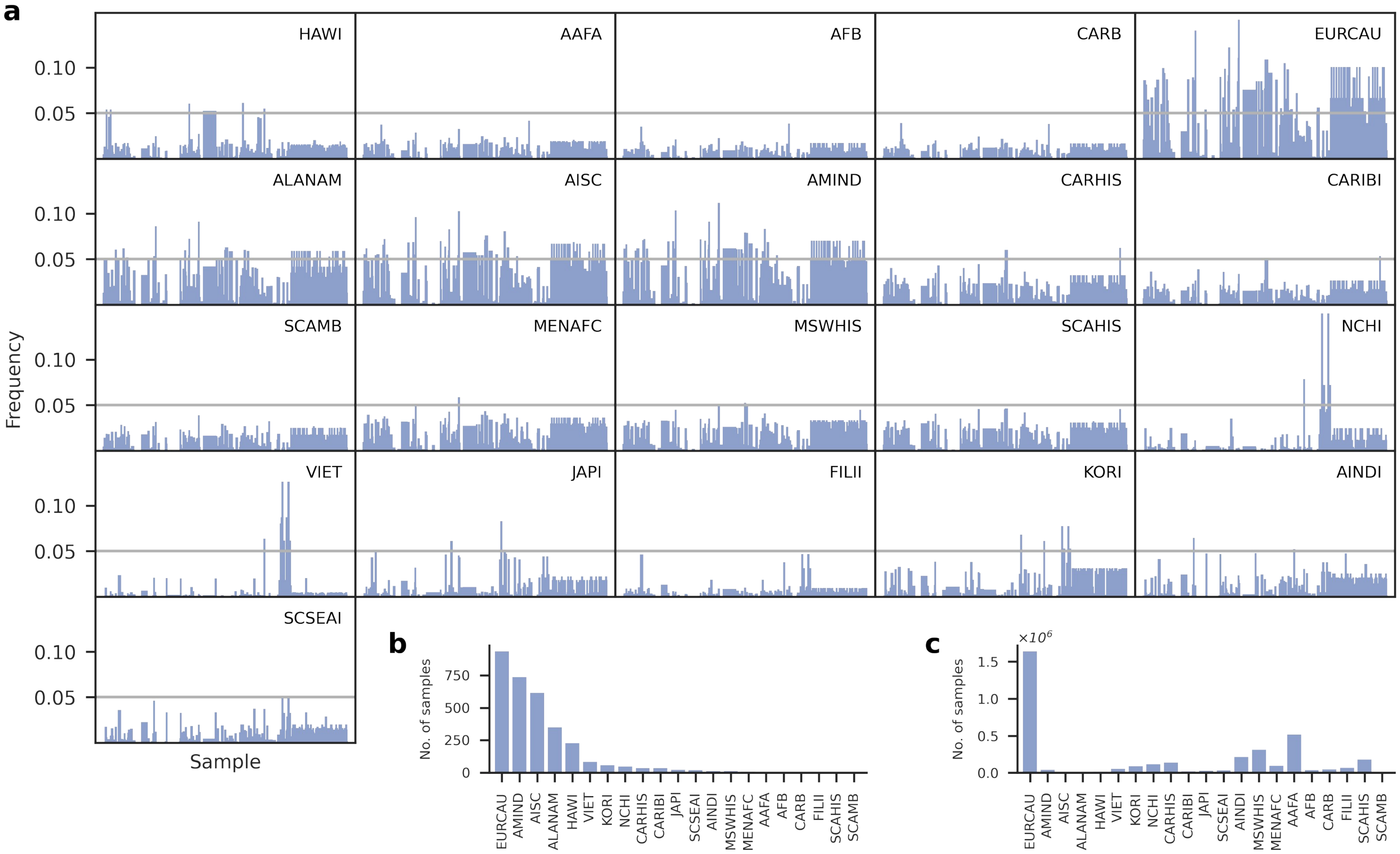


**Supplementary Figure 2: a)** The frequency distribution of the major HLA alleles shows similarities between most populations represented in the NMDP Haplotype Database except for certain East Asian communities such as VIET, which are distinct from other groups. An analysis of the number of samples belonging to each group revealed that Caucasian groups (EURCAU) are more represented in the database than others: **(b)** A frequency distribution of the sample numbers and **(c)** A scaled representation of the same. This emphasizes the need for greater diversity in the representation of population haplotype frequencies.
