## Supplementary material for "Expanding the definition of MHC Class I peptide binding promiscuity to support vaccine discovery across cancers with CARMEN": Supplementary File 1.pdf

**Supplementary File 1: MS-GF+ search parameters used to perform the closed immunopeptidomic database search**

```
#SpectrumFile
#  *.mzML, *.mzXML, *.mgf, *.ms2, *.pkl or *_dta.txt
#  Spectra should be centroided (see below for MSConvert example). Profile spectra will be ignored.
#  Use of -s at the command line will override this filename
#SpectrumFile=InstrumentFile.mzML

#FASTA file
#  "*.fasta or *.fa or *.faa
#  Use of -d at the command line will override this filename
#DatabaseFile=Proteins.fasta

#Prefix for decoy proteins in the FASTA file
#DecoyPrefix=XXX

#Precursor mass tolerance
#  Examples: 2.5Da or 30ppm
#  Use comma to set asymmetric values, for example "0.5Da,2.5Da" will set 0.5Da to the left (expMass<theoMass) and 2.5Da to the right (expMass>theoMass)
PrecursorMassTolerance=20ppm

#Max Number of Dynamic (Variable) Modifications per peptide
#  Default: 3
#  If this value is large, the search will be slow
NumMods=3

#Modifications (see below for examples)
#StaticMod=C2H3N1O1,     C,  fix, any,       Carbamidomethyl       # Fixed Carbamidomethyl C (alkylation)
#StaticMod=229.1629,     *,  fix, N-term,    TMT6plex
#StaticMod=229.1629,     K,  fix, any,       TMT6plex

# no variable modification should be added for first round search
# the pipeline with automatically add the 10 most abundant PTMs found by the open search
# DynamicMod=C2H2O,     *,  opt, N-term,      Acetyl             # Acetylation N-term
# DynamicMod=O1,        M,  opt, any,         Oxidation          # Oxidation M
# DynamicMod=HO3P,      STY,opt, any,         Phospho            # Phosphorylation STY


#Custom AA specification
```

CustomAA=C3H5NO,        U, custom, U, Selenocysteine   # Custom amino acids can only have C, H, N, O, and S
#CustomAA=C6H11NO,       X, custom, X, Leu_Ile         # Leucine or Isoleucine

#Fragmentation Method
###  0 means as written in the spectrum or CID if no info (Default)
###  1 means CID
###  2 means ETD
###  3 means HCD
###  Disabled in MS-GF+: 4 means Merge spectra from the same precursor (e.g. CID/ETD pairs, CID/HCD/ETD triplets)
FragmentationMethodID=0

#Instrument ID
###  0 means Low-res LCQ/LTQ (Default for CID and ETD); use InstrumentID=0 if analyzing a dataset with low-res CID and high-res HCD spectra
###  1 means High-res LTQ (Default for HCD; also appropriate for high res CID).  Do not merge spectra (FragMethod=4) when InstrumentID is 1; scores will degrade
###  2 means TOF
###  3 means Q-Exactive
InstrumentID=3

#Enzyme ID
###  0 means No enzyme used
###  1 means Trypsin (Default); use this along with NTT=0 for a no-enzyme search of a tryptically digested sample
###  2: Chymotrypsin, 3: Lys-C, 4: Lys-N, 5: Glu-C, 6: Arg-C, 7: Asp-N, 8: alphaLP, 9: No Enzyme (for peptidomics)
EnzymeID=0

#Isotope error range
###  Takes into account of the error introduced by choosing non-monoisotopic peak for fragmentation.
###  Useful for accurate precursor ion masses
###  Ignored if the parent mass tolerance is > 0.5Da or 500ppm
###  The combination of -t and -ti determins the precursor mass tolerance.
###  e.g. "-t 20ppm -ti -1,2" tests abs(exp-calc-n*1.00335Da)<20ppm for n=-1, 0, 1, 2.
IsotopeErrorRange=-1,2

#Number of tolerable termini
###  The number of peptide termini that must have been cleaved by the enzyme (default 1)
###  For trypsin, 2 means fully tryptic only, 1 means partially tryptic, and 0 means no-enzyme search

```
NTT=0

#Target/Decoy search mode
#  0 means don't search decoy database (default)
#  1 means search decoy database to compute FDR (source FASTA file must be forward-only proteins)
TDA=0

#Number of concurrent threads to be executed
#  Default: Number of available cores
#  To use three threads use NumThreads=3
NumThreads=All

#Minimum peptide length to consider
#  Default: 6
MinPepLength=8

#Maximum peptide length to consider
#  Default: 40
MaxPepLength=25

#Minimum precursor charge to consider (if not specified in the spectrum file)
#  Default: 2
MinCharge=1

#Maximum precursor charge to consider (if not specified in the spectrum file)
#  Default: 3
MaxCharge=4

#Number of matches per spectrum to be reported
#  If this value is greater than 1, the FDR values computed by MS-GF+ will be skewed by high-scoring 2nd and 3rd hits
NumMatchesPerSpec=1

#Mass of charge carrier
#  Default: mass of proton
#ChargeCarrierMass=1.00727649

#Maximum missed cleavages
#  Exclude peptides with more than this number of missed cleavages from the search, Default: -1 (no limit)
#MaxMissedCleavages=-1
```

#Minimum number of peaks per spectrum, Default:
###  Default: 10
MinNumPeaksPerSpectrum=10

#Number of isoforms to consider per peptide
###  Default: 128
#NumIsoforms=128

#Amino Acid Modification Examples
### Specify static modifications using one or more StaticMod= entries
### Specify dynamic modifications using one or more DynamicMod= entries
### Modification format is:
### Mass or CompositionString, Residues, ModType, Position, Name (all the five fields are required).
### CompositionString can only contain a limited set of elements, primarily C H N O S or P

### Examples:
#   C2H3N1O1,  C,  fix, any,         Carbamidomethyl    # Fixed Carbamidomethyl C (alkylation)
#   15.994915, M,  opt, any,         Oxidation          # Oxidation M (mass is used instead of CompositionString)
#   H-1N-1O1,  NQ, opt, any,         Deamidated         # Negative numbers are allowed.
#   CH2,       K,  opt, any,         Methyl             # Methylation K
#   C2H2O1,    K,  opt, any,         Acetyl             # Acetylation K
#   C2H3NO,    *,  opt, N-term,      Carbamidomethyl    # Variable Carbamidomethyl N-term
#   H-2O-1,    E,  opt, N-term,      Glu->pyro-Glu      # Pyro-glu from E
#   H-3N-1,    Q,  opt, N-term,      Gln->pyro-Glu      # Pyro-glu from Q
