## Supplementary material for "Expanding the definition of MHC Class I peptide binding promiscuity to support vaccine discovery across cancers with CARMEN": Supplementary File 2.pdf

**Supplementary File 2: MSFragger search parameters used to perform the open immunopeptidomic database search**

database_name = database.fasta # Path to the protein database file in FASTA format.
num_threads = 0 # Number of CPU threads to use.

precursor_mass_lower = -300 # Lower bound of the precursor mass window.
precursor_mass_upper = 300 # Upper bound of the precursor mass window.
precursor_mass_units = 0 # Precursor mass tolerance units (0 for Da, 1 for ppm).
precursor_true_tolerance = 20 # True precursor mass tolerance (window is +/- this value).
precursor_true_units = 1 # True precursor mass tolerance units (0 for Da, 1 for ppm).
fragment_mass_tolerance = 20 # Fragment mass tolerance (window is +/- this value).
fragment_mass_units = 1 # Fragment mass tolerance units (0 for Da, 1 for ppm).
calibrate_mass = 1 # Perform mass calibration (0 for OFF, 1 for ON, 2 for ON and find optimal parameters).
write_calibrated_mgf = 1 # Write calibrated MS2 scan to a MGF file (0 for No, 1 for Yes).
decoy_prefix = rev_ # Prefix added to the decoy protein ID.

isotope_error = 0 # Isotope correction for MS/MS events triggered on isotopic peaks.
mass_offsets = 0 # Creates multiple precursor tolerance windows with specified mass offsets.
precursor_mass_mode = selected # One of isolated/selected/recalculated.

localize_delta_mass = 1 # This allows shifted fragment ions - fragment ions with mass increased by the calculated mass difference,
### to be included in scoring.
delta_mass_exclude_ranges = (-1.5,3.5) # Exclude mass range for shifted ions searching.
fragment_ion_series = b,y # Ion series used in search.

search_enzyme_name = nonspecific # Name of enzyme to be written to the pepXML file.
search_enzyme_cutafter = ARNDCQEGHILKMFPSTWYV # Residues after which the enzyme cuts.
search_enzyme_butnotafter = # Residues that the enzyme will not cut before.

num_enzyme_termini = 0 # 0 for non-enzymatic, 1 for semi-enzymatic, and 2 for fully-enzymatic.
allowed_missed_cleavage = 25 # Allowed number of missed cleavages.

clip_nTerm_M = 1 # Specifies the trimming of a protein N-terminal methionine as a variable modification (0 or 1).

### maximum of 7 mods - amino acid codes, * for any amino acid,
### [ and ] specifies protein termini, n and c specifies
### peptide termini
### variable_mod_01 = 15.99490 M
### variable_mod_02 = 79.96633 STY
### variable_mod_03 = 42.01060 [^
### variable_mod_04 = -17.02650 nQnC
### variable_mod_05 = -18.01060 nE
### variable_mod_06 = 0.00000 site_06
### variable_mod_07 = 0.00000 site_07

allow_multiple_variable_mods_on_residue = 0 # Allow each amino acid to be modified by multiple variable modifications (0 or 1).
max_variable_mods_per_mod = 3 # Maximum number of residues that can be occupied by each variable modification (maximum of 5).
max_variable_mods_combinations = 5000 # Maximum allowed number of modified variably modified peptides from each peptide sequence, (maximum of 65534).

output_file_extension = pepXML # File extension of output files.
output_format = pepXML # File format of output files (pepXML or tsv).
output_report_topN = 1 # Reports top N PSMs per input spectrum.
output_max_expect = 50 # Suppresses reporting of PSM if top hit has expectation greater than this threshold.
report_alternative_proteins = 0 # Report alternative proteins for peptides that are found in multiple proteins (0 for no, 1 for yes).

precursor_charge = 1 4 # Assume range of potential precursor charge states. Only relevant when override_charge is set to 1.
override_charge = 0 # Ignores precursor charge and uses charge state specified in precursor_charge range (0 or 1).

digest_min_length = 8 # Minimum length of peptides to be generated during in-silico digestion.
digest_max_length = 25 # Maximum length of peptides to be generated during in-silico digestion.
digest_mass_range = 300.0 5000.0 # Mass range of peptides to be generated during in-silico digestion in Daltons.
max_fragment_charge = 2 # Maximum charge state for theoretical fragments to match (1-4).

# excluded_scan_list_file =                # If providing a text file containing a list of scan names, those scans would
### be ignored in the searching.

track_zero_topN = 0                # Track top N unmodified peptide results separately from main results internally for boosting features. Should be
### set to a number greater than output_report_topN if zero bin boosting is desired.
zero_bin_accept_expect = 0.00              # Ranks a zero-bin hit above all non-zero-bin hit if it has expectation less than this value.
zero_bin_mult_expect = 1.00              # Multiplies expect value of PSMs in the zero-bin during results ordering (set to less than 1 for boosting).
add_topN_complementary = 0              # Inserts complementary ions corresponding to the top N most intense fragments in each experimental spectra.

minimum_peaks = 15                # Minimum number of peaks in experimental spectrum for matching.
use_topN_peaks = 150                # Pre-process experimental spectrum to only use top N peaks.
min_fragments_modelling = 2              # Minimum number of matched peaks in PSM for inclusion in statistical modeling.
min_matched_fragments = 4              # Minimum number of matched peaks for PSM to be reported.
minimum_ratio = 0.01                # Filters out all peaks in experimental spectrum less intense than this multiple of the base peak intensity.
clear_mz_range = 0.0 0.0              # Removes peaks in this m/z range prior to matching.
remove_precursor_peak = 0              # Remove precursor peaks from tandem mass spectra. 0 = not remove; 1 = remove the peak with precursor charge;
### 2 = remove the peaks with all charge states.
remove_precursor_range = -1.5,1.5          # m/z range in removing precursor peaks. Unit: Da.
intensity_transform = 0              # Transform peaks intensities with sqrt root. 0 = not transform; 1 = transform using sqrt root.

### Fixed modifications
add_Cterm_peptide = 0.000000
add_Nterm_peptide = 0.000000
add_Cterm_protein = 0.000000
add_Nterm_protein = 0.000000
add_G_glycine = 0.000000
add_A_alanine = 0.000000
add_S_serine = 0.000000
add_P_proline = 0.000000
add_V_valine = 0.000000

add_T_threonine = 0.000000
add_C_cysteine = 0.000000
add_L_leucine = 0.000000
add_I_isoleucine = 0.000000
add_N_asparagine = 0.000000
add_D_aspartic_acid = 0.000000
add_Q_glutamine = 0.000000
add_K_lysine = 0.000000
add_E_glutamic_acid = 0.000000
add_M_methionine = 0.000000
add_H_histidine = 0.000000
add_F_phenylalanine = 0.000000
add_R_arginine = 0.000000
add_Y_tyrosine = 0.000000
add_W_tryptophan = 0.000000
add_B_user_amino_acid = 0.000000
add_J_user_amino_acid = 0.000000
add_O_user_amino_acid = 0.000000
add_U_user_amino_acid = 0.000000
add_X_user_amino_acid = 0.000000
add_Z_user_amino_acid = 0.000000
