## Supplementary material for "Expanding the definition of MHC Class I peptide binding promiscuity to support vaccine discovery across cancers with CARMEN": Supplementary File 3.pdf

**Supplementary Methods**

### 1. Immunopeptidomics from the Genomic perspective: Mapping peptides into contiguous genomic regions (contigs) and scaffolding these to capture enriched regions of antigen presentation.

The human genome and proteome were annotated with regions indicating the observed MHC Class I peptide sequences identified and their chromosomal coordinates. Subsequently, we developed a data model centered on these genomic mappings.

#### 1.1 A proteogenomic data model for epitopes, epitope contigs and epitope scaffolds

We developed a hierarchical proteogenomic data model from the peptide sequences and their mapped genomic coordinates, with the objective of compiling statistics around antigen presentation in a manner interoperable with other tools used in the genomic community, for example Bedtools. This involved collapsing the genome-mapped peptides into contigs and scaffolds through a series of views from the core data tables and associating statistics with these. This way, the genomic regions across our 2324 samples can be quickly understood in terms of their frequency of observation. Our data model has four layers corresponding to these different views.

1) *Whole genome mapping view (layer 1)*: This layer describes the mapping of peptides to the genome and tracks the genomic region of each peptide, the number of unique peptides covering that region and the unique peptide spectrum matches (PSMs). Unique PSMs are similar to reads in genomics and unique peptides can be interpreted as unique-reads. This view is derived from table "**carmen-mapped-protein-annotations-pogo.parquet**" converting the table to a standard BED file format, followed by using the genomic coordinates in the BED file to compute both peptide and spectral "coverage" over genomic regions corresponding to mapped peptides with the "GRanges" package version 1.46.1 in R version 4.1.2. (**Note: Peptide level coverage refers to the number of unique overlapping peptides mapping over a certain genomic region whereas spectral level coverage refers to the number of unique overlapping spectra mapping over a certain genomic region.) We also include as columns chromosome-level data (chromosome number, start and end) and the supporting peptide level data, including coverage, that correspond to their respective regions with START and END chromosomal coordinates.

2) *Clusters of unique peptides view (Layer 2)*: This view is identical to layer 1, but protein focused and includes protein coordinate information for each peptide. This is for the purposes of deriving protein-level hotspots as it is standard of practice in the field. This view can be used to associate the clustering of unique peptide sequences corresponding to a genomic region to the proteomic consequences of these clusterings. The number of unique peptides may be greater than or equal to one in any given region. These clusters represent peptides originating from the same genomic region with similar sequences.

This level is derived from the "**carmen-main.parquet**" and the "**carmen-mapped-protein-annotations-msfragger.parquet**" tables by combining the genomic coverage information obtained in the previous layer with the peptide-level sequence data taken from the main table mentioned above.

3) *Epitope contig view (Layer 3)*: This is a genome-centric view that collapses clusters of peptides to produce a higher level of organization known as "epitope contigs". These contigs represent "contiguous" stretches of peptide sequence clusters at a particular genomic location and are derived from the "**carmen-main.parquet**", "**carmen-mapped-protein-annotations-msfragger.parquet**" and the "**carmen-mapped-protein-annotations-pogo.parquet**" tables.

This view was derived as follows: The genomic distance between the overlapping peptides was measured and all peptides that occurred in contiguous regions were included into one "contig", whereas those falling outside this range were included in separate contigs. No distinction was made between peptides originating from "canonical" or "non-canonical" regions. The basic attributes of the epitope contig included chromosome number, starting and ending coordinates and a unique epitope contig identifier "Cx" (where x = 1..total number of contigs).

4) *Epitope scaffolds (Layer 4)*: Epitope contigs are separated by varying distances. The space between two contigs is called a "gap," and the "gap width" is defined as the distance measured by subtracting the ending coordinate of one contig from the starting coordinate of the other contig in the pair. Therefore, the gap width controls the number of contigs found within a scaffold. Scaffolds, representing the highest level of data organization, may consist of either a single contig or multiple contigs.

**2. Predicting antigen presentation with TransPHLA: model design, training, and testing**

We utilized TransPHLA, a published (https://www.nature.com/articles/s42256-022-00459-7) predictive model that captures the complex interactions between peptides and HLA class I molecules. Our goal was to understand the actual impact of including HLA Class I allele into the prediction of MHC Class I antigen presentation. To this end, we used the code-base for TransPHLA (https://github.com/a96123155/TransPHLA-AOMP), a transformer-based neural network designed to model these inter-residue dependencies with high fidelity. By leveraging the self-attention mechanism inherent to transformer architectures, TransPHLA considers both the peptide and HLA sequences. We addressed our question simply by adjusting the way the model was trained and tested, no further changes to the codebase altering how the model functioned were made.

The TransPHLA model was selected as one of the established and well-performing peptide-HLA binding predictors and a part of a wider vaccine development pipeline. This model is a transformer-based deep learning model with an attention mechanism developed to predict the binding affinity between peptides (8–14 amino acids long) and HLA class I molecules (up to 34 amino acids). The model consists of an embedding block that converts amino acids into numerical embeddings with added positional embeddings. Furthermore, an encoder block employs a masked multi-head self-attention mechanism, allowing the model to dynamically focus on the most relevant residues in the sequences. The peptide and HLA representations are concatenated to create a unified embedding of the peptide-HLA pair, which undergoes further encoding and feature optimization, and consequently is passed through a projection block to generate a predictive binding affinity score.

The aim was to investigate the importance of including the complete peptide-HLA pair sequences for antigen presentation. To evaluate this, we prepared six different ways of setting the training and testing data, and subsequently developed and tested models based on it. We used peptide and HLA class I molecule sequence datasets published by the authors of the TransPHLA model (for training, training validation, independent testing, and external testing) and modified it by randomly shuffling specific subsets of sequences within each corresponding file, which resulted in the following versions of input data:

(1) **Training HLA randomized:** HLA sequences shuffled in the training and validation datasets only
(2) **Testing HLA randomized:** HLA sequences shuffled in the independent and external datasets only
(3) **All HLA randomized:** HLA sequences shuffled in all dataset files
(4) **Training peptides randomized:** peptide sequences shuffled in the training and validation datasets only
(5) **Testing peptides randomized:** peptide sequences shuffled in the independent and external datasets only
(6) **All peptides randomized:** peptide sequences shuffled in all dataset files.

Because of some overlap between peptide and HLA sequences in the original datasets, the shuffling process prevented creating duplicate peptide-HLA pairs across files, which could skew results. Once a peptide-HLA pair was assigned to one file, it was excluded from all others.

After generating the input data, we conducted a series of tests to evaluate the model. Using a model development pipeline provided by the authors (“pHLAIformer.ipynb”), we trained and assessed the core model’s ability to classify peptides for the aforementioned cases and compared its results with unchanged original published data (Source dataset). **Figure 3d** is included below for context.

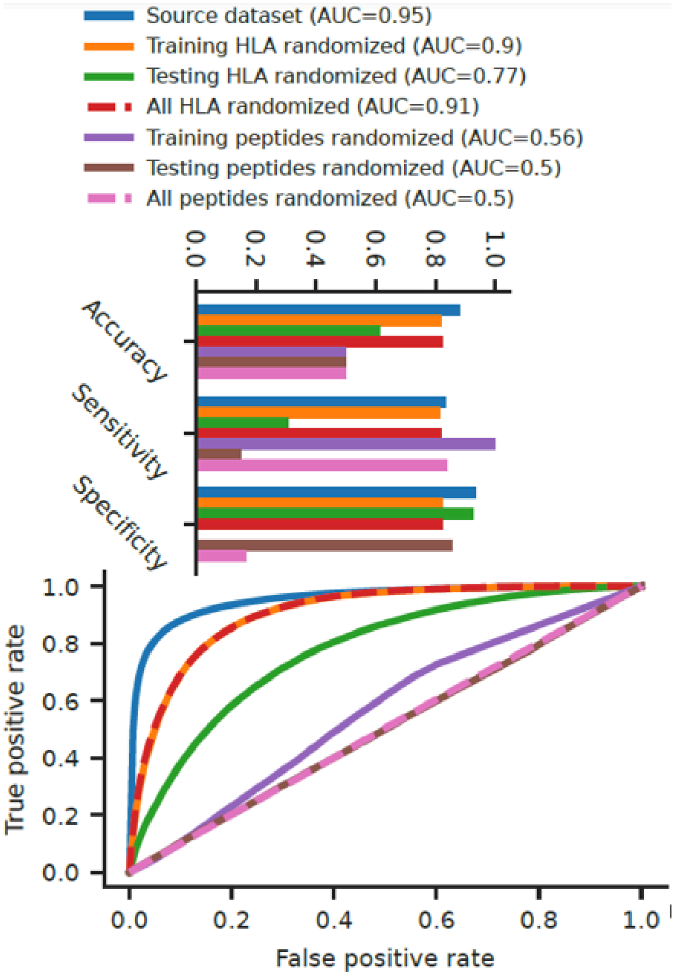


**Figure 3d**

**3. Predicting Immunotherapy response with Support Vector Machines: SVM model**

Finally, we explored the utility of understanding whether a mutation that either overlapped a peptide previously observed in the immunopeptidome, a contig or scaffolds, alongside other metrics like promiscuity associated with those genomic regions was useful for predicting response to immunotherapy leveraging the CARMEN database.

To explore this research question, we utilized a mutation database associated with the Braun dataset (Braun et al. 2020). The full dataset contains 311 patients and integrated molecular, clinical, and tumor-specific features relevant to treatment response in metastatic renal cell carcinoma. Columns in the table were grouped when considered in the SVM model as follows.

Molecular Variables:

- GE: normalized gene expression values, reduced using PCA down to 200 dimensions.

- BP: tumor-level pathway enrichment scores computed using single-sample Gene Set Enrichment Analysis (ssGSEA) across Reactome, KEGG, PID, and BIOCARTA biological pathway databases, reduced using PCA down to 100 dimensions,
- TF: tumor mutation burden (TMB), immunofluorescence-derived CD8+ T cell infiltration scores at both the tumor core and periphery, as well as tumor purity and ploidy estimates.
- MT: The chromosomal position of a mutation, either alone or augmented with additional immunopeptidomic features associated with the mutation region, reduced to 4000 dimensions via random indexing as described below this paragraph.

Clinical Variables

- PS: MSKCC and IMDC prognostic risk scores
- CF: sex and age, as well as the information on whether the patient received prior therapies. Also the arm of clinical trial, clinical benefit, ORR, PFS, OS.

The tables are available in the git-repository (https://github.com/immuno-informatics/SVM_Immunotherapy/tree/main/data).

All features were selected based on their biological relevance and potential associations with drug response. These were developed into a table containing prognostic scores (PS), biological processes (BP), tumor/clinical features (TF/CF), mutations (MT), and gene expression (GE), were evaluated.

We aimed to investigate whether a mutation in an epitope scaffold may be more predictive for the response to treatment concerning the overall mutations. Given that the Braun dataset is small compared to what is needed for a neural network, we used a variety of machine learning models, and we reported results based on Support Vector Machines (SVM) with three different kernels: a linear kernel, a radial basis function kernel and a polynomial kernel with a degree of 2.

To enable the exploration of the research question by using SVM, we used Random Indexing to accommodate the feature space of gene mutations within a reasonable size feature space. In Random Indexing, each one-hot feature is represented as a random vector drawn from a multinomial normal distribution. Given a patient P with a set of relevant mutations X, we compute the sum of the vectors representing the mutations in X, and then we associate this vector sum with the patient P.

Using Random Indexing of mutations and two other groups of features for the clinical history and the gene expression of the patient, we had the possibility to experiment with many configurations, which vary on the way that mutation information is extended:

- Peptide Level -- peptides selected, number of unique HLAs and promiscuity calculated as the set of peptides directly overlapping the mutation
- Contig Level -- peptides selected, number of unique HLAs and promiscuity calculated for the set of peptides in contig

- Scaffold Level -- peptides selected, number of unique HLAs and promiscuity calculated for the set of peptides in scaffold
- Baseline -- mapping of immunopeptidomics to genomics not considered

For each of the above cases an optimization in the space of combinations of additional information were examined, additionally allowing for scaling each group of features by a factor between 0.0 and 1.0 with a granularity of 0.1. Utilizing the Optuna library, each case was allowed at least 10,000 random samples, with an additional 10,000 with fixed parameters of radial basis function kernel and the inclusion of promiscuity as extended mutation information - for the cases including immunopeptidomics information. Radial basis function kernel and the inclusion of promiscuity were chosen as fixed for consistently being among the top performing sets in the first 10,000 samples.
